# Amino acid transporter SLC7A5 regulates Paneth cell function to affect the intestinal inflammatory response

**DOI:** 10.1101/2023.01.24.524966

**Authors:** Lingyu Bao, Liezhen Fu, Yijun Su, Zuojia Chen, Zhaoyi Peng, Lulu Sun, Frank J. Gonzalez, Chuan Wu, Hongen Zhang, Bingyin Shi, Yun-Bo Shi

## Abstract

The intestine is critical for not only processing and resorbing nutrients but also protecting the organism from the environment. These functions are mainly carried out by the epithelium, which is constantly being self-renewed. Many genes and pathways can influence intestinal epithelial cell proliferation. Among them is mTORC1, whose activation increases cell proliferation. Here, we report the first intestinal epithelial cell-specific knockout (^ΔIEC^) of an amino acid transporter capable of activating mTORC1. We show that the transporter, SLC7A5, is highly expressed in mouse intestinal crypt and *Slc7a5*^ΔIEC^ reduces mTORC1 signaling. Surprisingly, *Slc7a5*^ΔIEC^ mice have increased cell proliferation but reduced secretory cells, particularly mature Paneth cells. scRNA-seq and electron microscopic analyses revealed dedifferentiation of Paneth cells in *Slc7a5*^ΔIEC^ mice, leading to markedly reduced secretory granules with little effect on Paneth cell number. We further show that *Slc7a5*^ΔIEC^ mice are prone to experimental colitis. Thus, SLC7A5 regulates secretory cell differentiation to affect stem cell niche and/or inflammatory response to regulate cell proliferation.

## Introduction

The intestine is the organ responsible for the digestion and absorption of nutrients and water, and also forms a barrier against environmental harmful materials/organisms. It is under a constant barrage of mechanical, chemical, and pathogen-driven attacks from food digestion and the trillions of microorganisms that colonize the intestine (*1, 2*). It is thus critical to maintain intestinal integrity and function. Intestinal homeostasis relies on a perfect balance of the interactions among different cell types within intestinal epithelium, the underlying non-epithelial tissues, the microbiota, and immune system (*2-5*). Intestinal epithelial cells are crucial for intestinal homeostasis and are constantly renewed, every 3-5 days in the case of mice, driven by adult stem cells (*6, 7*). Two types of epithelial cells, goblet cells and Paneth cells, function to maintain intestinal integrity. Goblet cells secret mucus that coats the epithelium to block the access of the microbiota to mucosa and to allow the accumulation of high concentrations of anti-microbial peptides (AMPs). Reduced goblet cell numbers and mucus secretion are the hall marks of human inflammatory bowel disease (IBD) and experimental dextran sulfate sodium (DSS)-induced colitis(*8*). Paneth cells secrete AMPs, including lysozyme and defensins, that protect the epithelium from pathogens. Recent studies have shown that impaired AMP secretion may contribute to IBD susceptibility(*9, 10*).

SLC7A5, also known as LAT1, forms a heterodimeric amino acid transporter by interacting with glycoprotein CD98 (SLC3A2) involving a conserved disulfide bond (*11*). It transports large neutral amino acids and is crucial for immune system, placenta, blood brain barrier, and protecting the body against pathogens (*12-14*). SLC7A5-mediated mTORC1 activation is well known to be important for immune system, especially T cell activation (*15*). SLC7A5 is broadly expressed in many organs, including the intestine (*16*). However, if and how SLC7A5-mediated mTORC1 activation plays a role in intestinal homeostasis and function remain unknown. While there have been studies on mTORC1 signaling in intestine, they have been mainly focused on energy utilization with not very consistent findings. For example, it was shown that calorie restriction leads to inhibition of mTORC1 pathway specifically in Paneth cells, which in turn increases crypt cell proliferation and LGR5+ stem cell numbers through activating SIRT1 (*17-19*). On the other hand, a different study showed that calorie restriction reduced cell proliferation, increased reserve stem cells (BMI1+ stem cells) other than LGR5+ stem cells in the intestine (*20*). Thus, further analyses are needed to understand the effects and regulation of mTORC1 signaling in intestinal homeostasis.

Here, we made use of a floxed *Slc7a5* allele (*Slc7a5*^fl/fl^) (*16*) to knock out *Slc7a5* specifically in the intestinal epithelium by crossing the *Slc7a5*^fl/fl^ line with a villin-cre line, where the Cre recombinase is under the control of intestinal epithelium-specific villin promoter, to generate *Slc7a5*^ΔIEC^ mice. This allowed us to investigate if epithelial expression of this amino acid transporter affects mTORC1 function and/or intestinal development and/or homeostasis. Our study showed that loss of SLC7A5 in intestinal epithelium led to reduced mTORC1 signaling and a dramatic decrease in the number of mature secretory cells, including Paneth cells and goblet cells. Consistent with the fact that these cells are the primary defense cells in intestine through establishment of the mucus layer and secreting AMPs, the *Slc7a5*^ΔIEC^ mice were prone to DSS-induced colitis. In addition, as mTORC1 is a well-known, important pathway for cell proliferation, we examined cell proliferation in the intestine and found that surprisingly, loss of SLC7A5 in intestinal epithelium increased cell proliferation in the crypts. Furthermore, we observed increased cell death in the epithelium and faster epithelial cell turnover. On the other hand, the number of crypt base columnar (CBC) stem cells, such as LGR5 positive and OLFM4 positive stem cells, were not changed. Our findings thus revealed that knocking out SLC7A5 in intestinal epithelium affected intestinal defense function via reduced mTORC1 signaling in secretory cell lineage and further suggest a model where a complex and delicate intestinal regulatory system helps to maintain physiological or pathophysiological status of the intestine.

## Results

### Intestinal epithelium-specific knockout of SLC7A5 leads to longer crypts in mice

Whole body knockout of SLC7A5 causes embryonic lethality in mice (*21*). To study the role of SLC7A5 in the intestine, we crossed villin-Cre mice with *Slc7a5*^fl/fl^ mice to delete the floxed exon in *Slc7a5* gene specifically in intestinal epithelium (Fig. 1A). Homozygous *Slc7a5*^ΔIEC^ mice were born at the Mendelian ratios and developed normally. Western blot analysis of isolated small intestinal crypts showed that SLC7A5 protein was expressed in the epithelium of wild-type *Slc7a5*^fl/fl^ mice but absent in *Slc7a5*^ΔIEC^ mice (Fig. 1B), demonstrating the knockout efficiency. Furthermore, immunohistochemical analysis showed that in adult intestine, SLC7A5 was found in both epithelium and submucosal area of wild-type mice but absent in the epithelium of the *Slc7a5*^ΔIEC^ mice (Fig. 1C). The *Slc7a5*^ΔIEC^ mice were phenotypically normal externally throughout development and adulthood (not shown). In addition, isolated intestine also had normal gross morphology (not shown). On the other hand, H&E-stained histological sections of the small intestine showed that the *Slc7a5*^ΔIEC^ intestine had longer crypts (Fig. 1D).

### Slc7a5^ΔIEC^ mice have increased cell proliferation in the crypt and accelerated epithelial cell turn-over with compensating increases in cell death in the epithelium

Generally, the upper two thirds of the crypt is referred to as the transit amplifying cell zone, where the cells, called transit amplifying cells, proliferate rapidly and therefore can be visualized with Ki67 labeling (Fig. 2A). In *Slc7a5*^ΔIEC^ mice, the Ki67 positive cells increased by about two-fold compared to that in *Slc7a5*^fl/fl^ mice (Fig. 2B). In addition, while rare, Ki67-positive proliferating cells could also be detected at the crypt base, intercalating among Paneth cells. Crypt base stem cell proliferation was also increased by *Slc7a5*^ΔIEC^ (Fig. 2C). On the other hand, *in situ* hybridization with probes for stem cell markers LGR5 and OLFM4 mRNAs showed no difference between *Slc7a5*^ΔIEC^ and *Slc7a5*^fl/fl^ mice (Supplementary Fig. 1).

Since intestinal gross morphology did not change very significantly, the increased cell proliferation in *Slc7a5*^ΔIEC^ mice suggest faster cell turn-over in the epithelium in order to maintain homeostasis. To test this, we carried out kinetic studies on cell turn-over by labeling proliferating cells with a single injection of Edu, followed by euthanizing the mice at 2 h, 8 h, 24 h, 48 h and 72 h post injection (Fig. 2D). Analysis of the Edu-labeled cells in the resulting intestinal cross-sections showed that at 2 and 8 h post injection, the Edu-positive cells per crypt in *Slc7a5*^ΔIEC^ mice were about two-folds of those in *Slc7a5*^fl/fl^ mice (Fig. 2E-F). Furthermore, at 8 h, many Edu-positive cells in *Slc7a5*^ΔIEC^ mice but few in *Slc7a5*^fl/fl^ mice had migrated out of crypts and into villi (Fig. 2E), suggesting faster migration and differentiation in knockout cells. By 24 h, most Edu-positive cells in *Slc7a5*^ΔIEC^ mice had migrated to near the top of the villi while most Edu-positive cells in *Slc7a5*^fl/fl^ mice were still near the lower half of the villi (Fig. 2E), indicating again that the *Slc7a5*^ΔIEC^ led to faster cell migration during epithelium turn-over. The Edu-positive cells reached maxima between 8-24 h and subsequently decreased as they turned over, with all removed by 48 h and 72 h in *Slc7a5*^ΔIEC^ and *Slc7a5*^fl/fl^ mice, respectively (Fig. 2E-F), demonstrating that *Slc7a5*^ΔIEC^ had faster turn-over. Consistently, TUNEL-labeling for apoptotic cells showed that apoptotic cells in small intestine were increased dramatically by *Slc7a5*^ΔIEC^ (Fig. 2G-H). Thus, *Slc7a5*^ΔIEC^ caused faster epithelium turn-over by increasing cell proliferation in the crypt with compensatory increases in cell migration and cell death.

**Figure 1.**
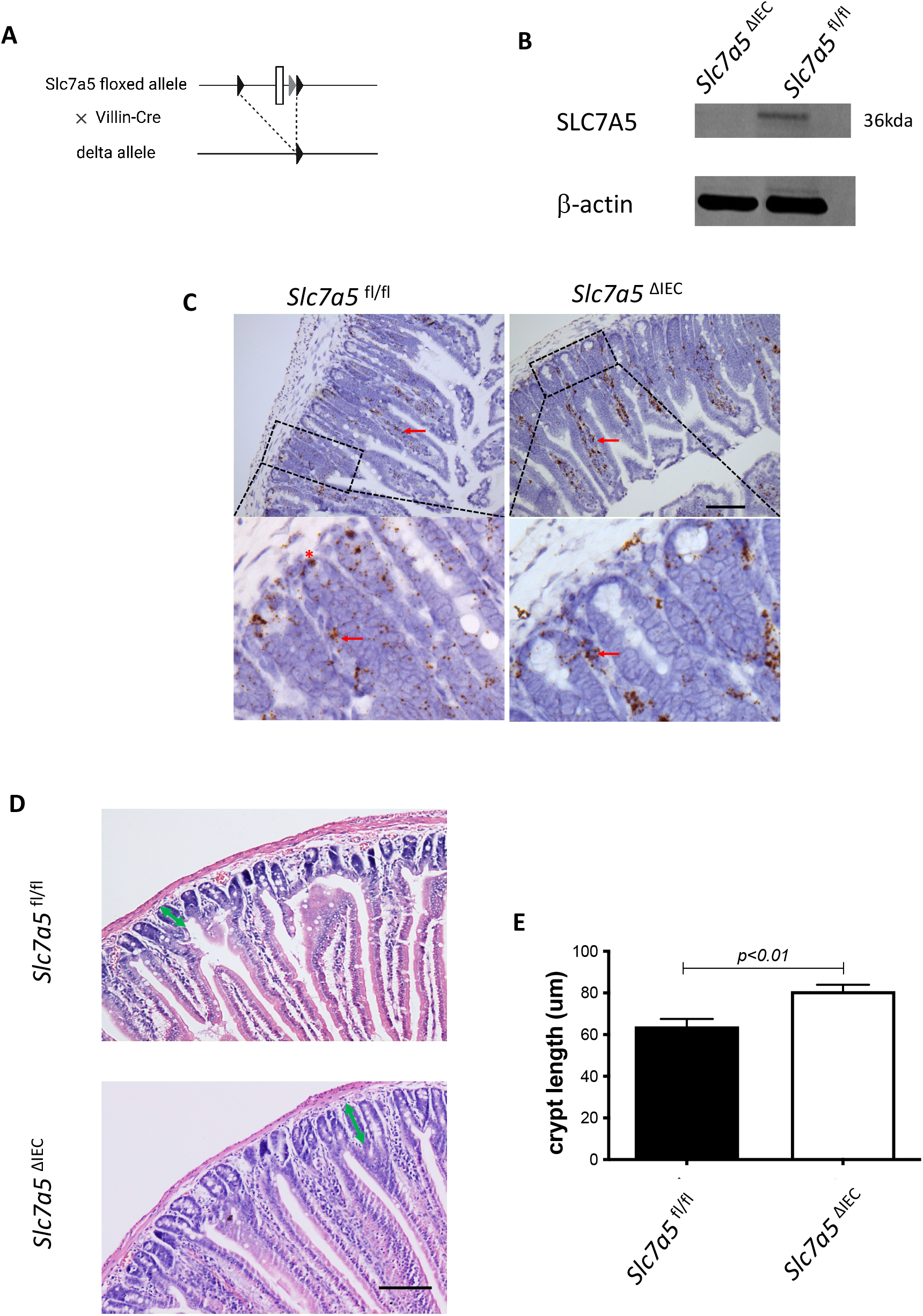
Intestinal epithelial cell specific knockout of SLC7A5 leads to more elongated crypts in the small intestine. **A**. Schematic diagram of the *Slc7a5*-floxed locus in *Slc7a5*^fl/fl^ mice (*16*) used to generate *Slc7a5*^ΔIEC^ mice. Two loxP sites (black triangle) flanked *Slc7a5* exon 1. When *Slc7a5*^fl/fl^ mice are crossed with mice containing Cre under the control of the epithelial specific villin promoter (Villin-Cre), the exon flanked by the two loxP sites is removed to generate the deleted allele. **B**. *Slc7a5*^ΔIEC^ mice have little or no detectable SLC7A5 protein in the intestinal epithelium. Intestinal epithelium was isolated from *Slc7a5*^ΔIEC^ and *Slc7a5*^fl/fl^ mice and its proteins were subjected to western blot analysis for *SLC7A5* expression, with β-actin as a loading control. **C**. SLC7A5 is expressed in both epithelium (red asterisk) and submucosal area (red arrows) and *Slc7a5*^ΔIEC^ eliminates its epithelial expression. Intestinal cross-sections were subjected to immunohistochemical analysis for SLC7A5 expression. Note that *Slc7a5*^ΔIEC^ reduced/abolished SLC7A5 in the crypt epithelium but not in the submucosal area, e.g., immune cells (red arrows). Scale bars, 100µm. **D-E**. *Slc7a5*^ΔIEC^ mice have longer crypts in the small intestine. HE staining of intestine from *Slc7a5*^fl/fl^ and *Slc7a5*^ΔIEC^ mice **D**. showing longer crypts (green double headed arrows). The crypt lengths were measured (n=5) and shown as mean ± SD (E).

**Figure 2.**
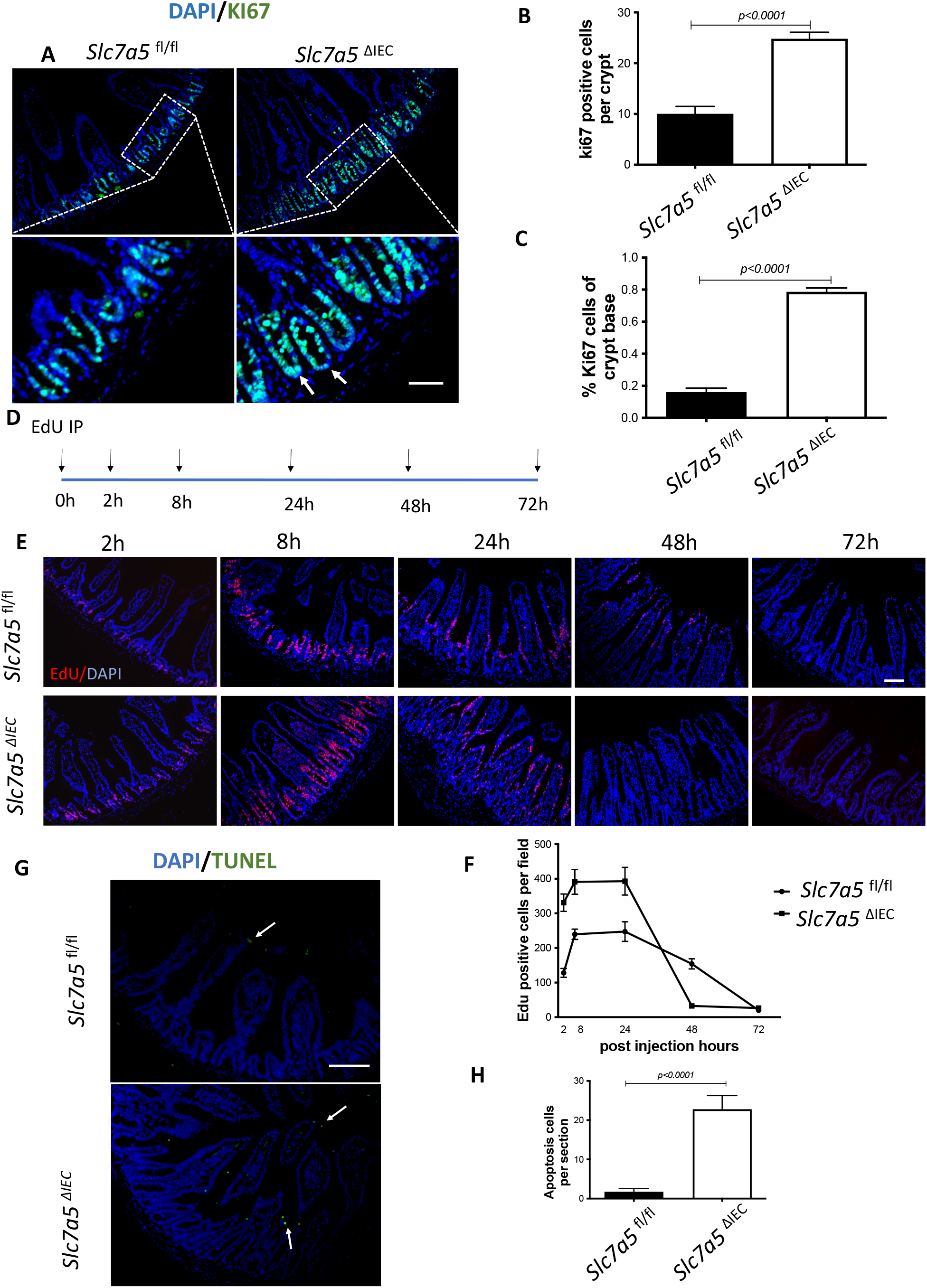
*Slc7a5*^ΔIEC^ mice have faster epithelial cell turnover, with increased cell proliferation and epithelial cell death. **A-C**. Cross-sections of the small intestine were labeled with DAPI (blue) for DNA and anti-Ki67 antibody (green) for proliferating cells (**A**), showing more cell proliferation in the crypt of *Slc7a5*^ΔIEC^ mice. The proliferating cells in the crypt (**B**) or at crypt base (**C**) were quantified, revealing that *Slc7a5*^ΔIEC^ mice had increased Ki67 staining in both transit amplifying (TA) cell zone and crypt base, where crypt base stem cells and Paneth cells are located, respectively. Scale bars, 100µm. **D**. Schematic diagram of Edu time course experiment. Edu was injected at 0 hr and the intestine was isolated at indicated time points (arrows) for analysis **E**. Edu staining of small intestine from *Slc7a5*^fl/fl^ and *Slc7a5*^ΔIEC^ mice at 2 h, 8 h, 24 h, 48 h and 72 h after Edu injection. Note that compared to *Slc7a5*^fl/fl^ mice, *Slc7a5*^ΔIEC^ mice had increased EdU-labeling in the crypts at 2-8 h, indicative of increased cell proliferation. The Edu-labeling reached plateau after 8 h, suggesting that the unincorporated Edu had been metabolized. The Edu-labeled cells in *Slc7a5*^ΔIEC^ intestine, however, had faster migration, with many of them differentiated and migrated into the villi by 8 h. Scale bars, 100µm. **F**. Quantification of Edu-labeled cells from intestinal cross-sections as shown in E. Note that *Slc7a5*^ΔIEC^ mice had increased cell proliferation (2-8 h) and faster turnover rate, with essentially all Edu-labeled cells eliminated by 48 h. G-H. TUNEL assay showing increased apoptotic cells in small intestine of *Slc7a5*^ΔIEC^ mice compared to *Slc7a5*^fl/fl^ mice (**G**), with the quantification shown mean ± SD (**H**). Scale bars, 100µm.

### Slc7a5^ΔIEC^ mice had fewer Paneth cells and goblet cells in the small intestine

We next analyzed if intestinal epithelial cell disruption of *Slc7a5* affected the Paneth cells and goblet cells, the main secretary cells in the small intestine. Immunohistochemical analysis with an antibody against lysozyme, a marker for mature Paneth cells, revealed that *Slc7a5*^ΔIEC^ mice had a dramatic reduction in number of mature Paneth cells compared to *Slc7a5*^fl/fl^ mice (Fig. 3A, B). Similarly, staining with Alcian blue for goblet cells revealed that the number of goblet cells were reduced in villi of *Slc7a5*^ΔIEC^ mice (Fig. 3C, D). Interestingly, more goblet cells were present in the crypts of *Slc7a5*^ΔIEC^ mice (Fig. 3C, E). Thus, *Slc7a5*^ΔIEC^ had abnormal development and/or distribution of the secretory cells.

**Figure 3.**
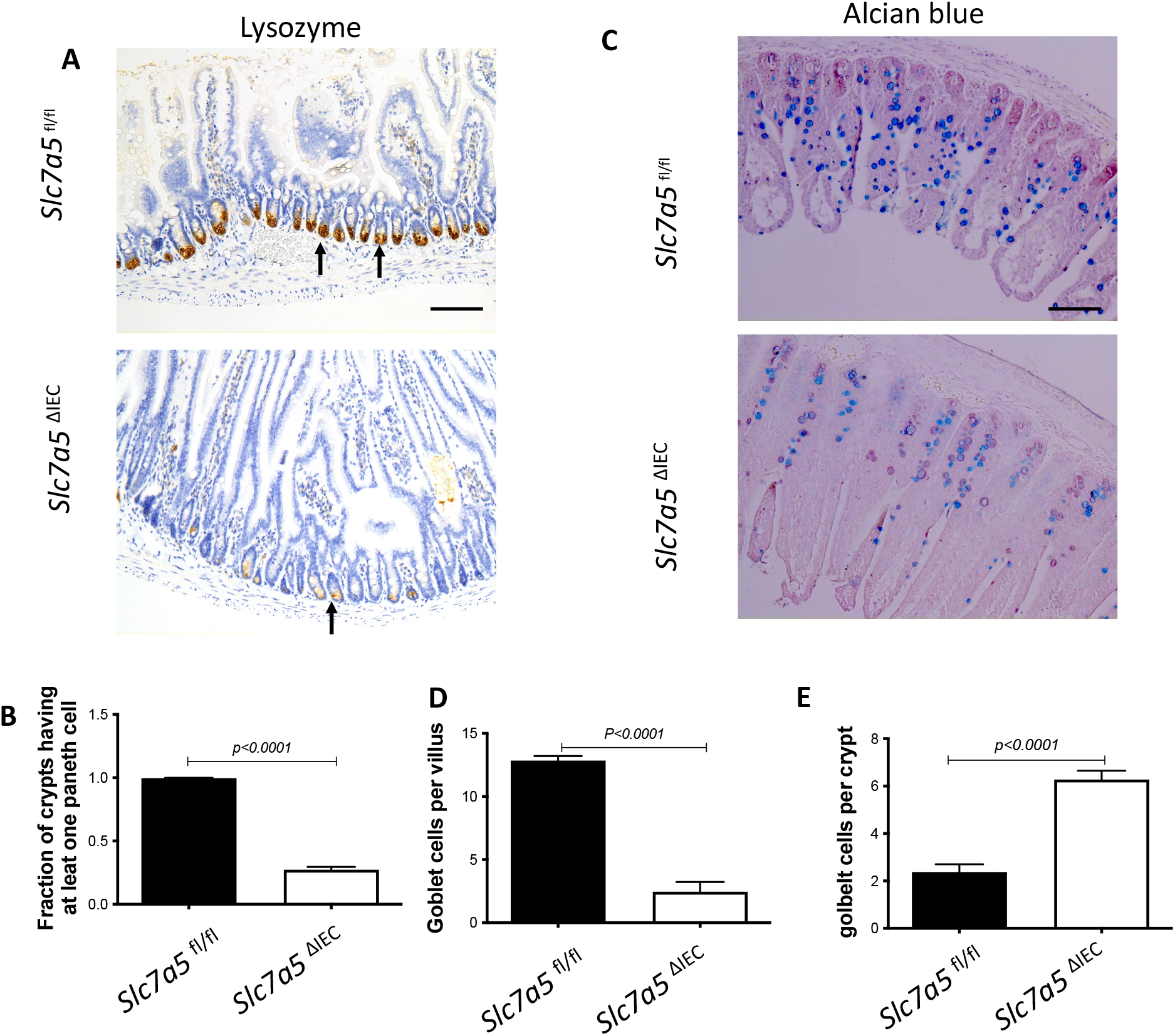
*Slc7a5*^ΔIEC^ causes defects in secretory cell lineages. **A-B**. Much fewer cells were stained with anti-lysozyme antibody for Paneth cells in small intestine sections of *Slc7a5*^ΔIEC^ mice compared to *Slc7a5*^fl/fl^ mice (A) with quantification of the number of lysozyme-positive Paneth cells/crypt shown in (**B**). Scale bars, 100µm. C-E. Alcian blue stained goblet cells (**C**) were reduced in villi (**D**) but increased in crypts (**E**) of small intestine of *Slc7a5*^ΔIEC^ mice compared to *Slc7a5*^fl/fl^ mice. Scale bars, 100µm. Quantifications were shown as mean ± SD.

### Slc7a5^ΔIEC^ mice have altered intestinal mTORC1 signaling

*Slc7a5* encodes SLC7A5, also called LAT1, which forms a heterodimeric transmembrane amino acid transporter with glycoprotein CD98 (SLC3A2). The heterodimer can transport L-branched chain amino acids and natural amino acids, leading to activation mTORC1 signaling. In addition, a mTORC1 and mTORC2 knockout study revealed reduced secretory cells (*22*). To investigate whether *Slc7a5*^ΔIEC^ reduces mTORC1 signaling to affect the secretory cell lineage, we analyzed the phosphorylation of mTORC1 substrates p70 ribosome protein S6 kinase1 (70S6K) and ribosome protein S6 (S6) (*23*). The results showed that the phosphorylated form of both 70S6K (i.e., p70S6K) (Fig. 4A, B) and S6 (i.e., pS6) (Fig. 4 C, D) was reduced in the small intestine of *Slc7a5*^ΔIEC^ mice compared to *Slc7a5*^fl/fl^ mice. To determine if the alteration in mTORC1 signaling was through amino acid transporter activity, we treated intestinal organoids made from intestinal crypts of *Slc7a5*^fl/fl^ and *Slc7a5*^ΔIEC^ mice with mTORC1 activator MHY1485 (*24*), mTORC1 inhibitor rapamycin (*24-26*), or amino acid transporter inhibitor BCH (*11, 27, 28*) (Fig. 4E, F). Consistent with the in *vivo* findings, *Slc7a5*^ΔIEC^ organoids had decreased pS6 level compared to *Slc7a5*^fl/fl^ mouse organoids under control condition. Rapamycin treatment of the *Slc7a5*^fl/fl^ mouse organoids led to expected inhibition of pS6. Importantly, MHY1485 treatment of *Slc7a5*^ΔIEC^ organoids increased pS6 to a similar level as in *Slc7a5*^fl/fl^ organoids, while BCH treatment of *Slc7a5*^fl/fl^ organoids reduced pS6 to a level similar to that in *Slc7a5*^ΔIEC^ mouse organoids. These findings not only showed that *Slc7a5*^ΔIEC^ reduced mTORC1 signaling through reduced amino acid transport but also suggest that SLC7A5 is the major amino acid transporter that can regulate mTORC1 signaling in the intestine.

**Figure 4.**
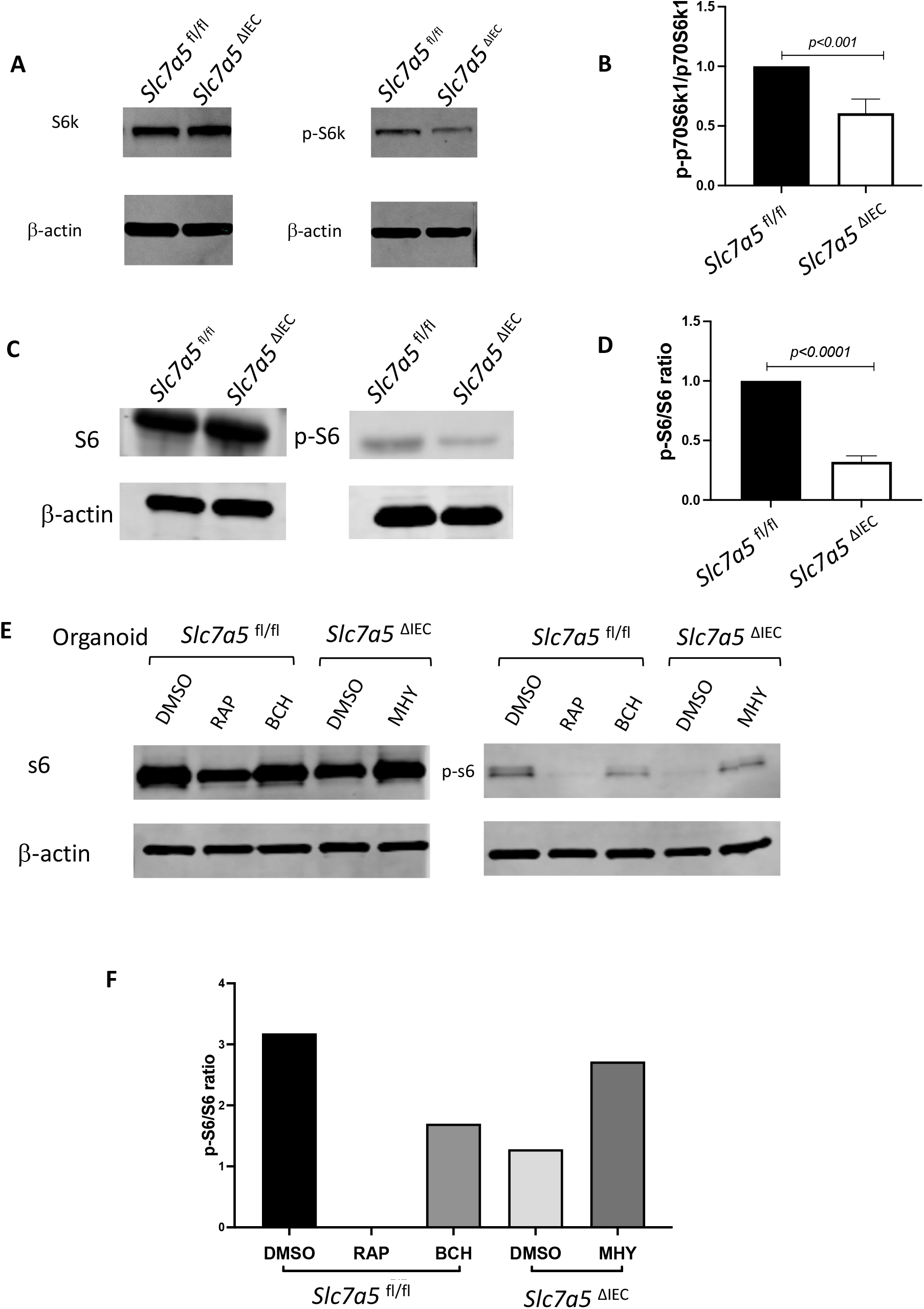
*Slc7a5*^ΔIEC^ leads to reduced mTORC1 signaling in intestinal epithelium. **A/C**. Western blot analysis of total S6k (**A**) and S6 (**C)**, downstream effectors of mTORC1 signaling, and phosphorylated S6k (p-S6k) (**A**) and phosphorylated S6 (p-S6) in intestinal epithelium; β-actin was used as a loading control. B/D. Quantification of the levels of P-S6k (**B**) and P-S6 **(D**), measures of mTORC1 signaling, from 5 (**B**) or 2 (**D**) independent experiments, revealed reduced mTORC1 signaling due to *Slc7a5*^ΔIEC^. **E-F**. *Slc7a5*^ΔIEC^ also reduces mTORC1 signaling in intestinal epithelial organoids *in vitro*. Small intestinal crypts from *Slc7a5*^fl/fl^ and *Slc7a5*^ΔIEC^ mice were cultured as organoids for totally 7 days. Some organoids were treated with BCH (10 mM) starting on day 4 for 3 days while some others were treated with rapamycin (20 µM) or MHY1485 (10 µM) on day 6 for one day, as indicated. On day 7, the organoids were isolated for western blot analysis for indicated proteins (**E**). The signals from the blots were quantified (**F**). Note the reduced p-S6 in *Slc7a5*^ΔIEC^ organoids compared to *Slc7a5*^fl/fl^ mouse organoids. As expected, treatment with MHY1485, an mTORC1 activator, increased p-S6 in *Slc7a5*^ΔIEC^ organoids while both and rapamycin, an mTORC1 inhibitor, and BCH, an SLC7A5 inhibitor, reduced p-S6 in *Slc7a5*^fl/fl^ mouse organoids, resembling *Slc7a5*^ΔIEC^ mice. Two independent experiments were done with similar results (not shown).

### Slc7a5^ΔIEC^ mice had Paneth dedifferentiation with reduced secretory granules

To further characterize the changes among different types of crypt cells caused by *Slc7a5*^ΔIEC^, we transcriptionally profiled single cells (single cell RNA-seq or scRNA-seq) from the crypts from the small intestine of *Slc7a5*^fl/fl^ and *Slc7a5*^ΔIEC^ mice. After unsupervised graph clustering, the sequenced cells were partitioned into different groups and the cell type identities of the groups were identified based on the expression of known marker genes(*29*) (Supplementary Fig. 2 and Supplementary Data 1). We found 6 known cell types in the *Slc7a5*^ΔIEC^ crypts and 4 in the *Slc7a5*^fl/fl^ crypts, and another group of unknown identity in the *Slc7a5*^ΔIEC^ sample (Fig. 5A and Supplementary Fig. 2). The two known cell types (enteroendocrine and Tuft cells) not found in the *Slc7a5*^fl/fl^ mouse crypts were likely due to fewer cells were sequenced in the *Slc7a5*^fl/fl^ sample compared to the *Slc7a5*^ΔIEC^ sample since they represented only a small fraction of the total cells (Fig. 5A). Quantification of the percentage of each cell type within the sample for each animal group showed that *Slc7a5*^ΔIEC^ led to increased goblet cell, increased proliferating cells (TA or cell cycle group), and reduced Paneth cells (Fig. 5A), all in agreement with the histology data above. In addition, the feature genes for the group of cells labeled as unclassified cells (the unknown group) were mostly mitochondria genes, which indicated that this group of cells were apoptotic cells, consistent with increased apoptosis in *Slc7a5*^ΔIEC^ intestine as shown above.

**Figure 5:**
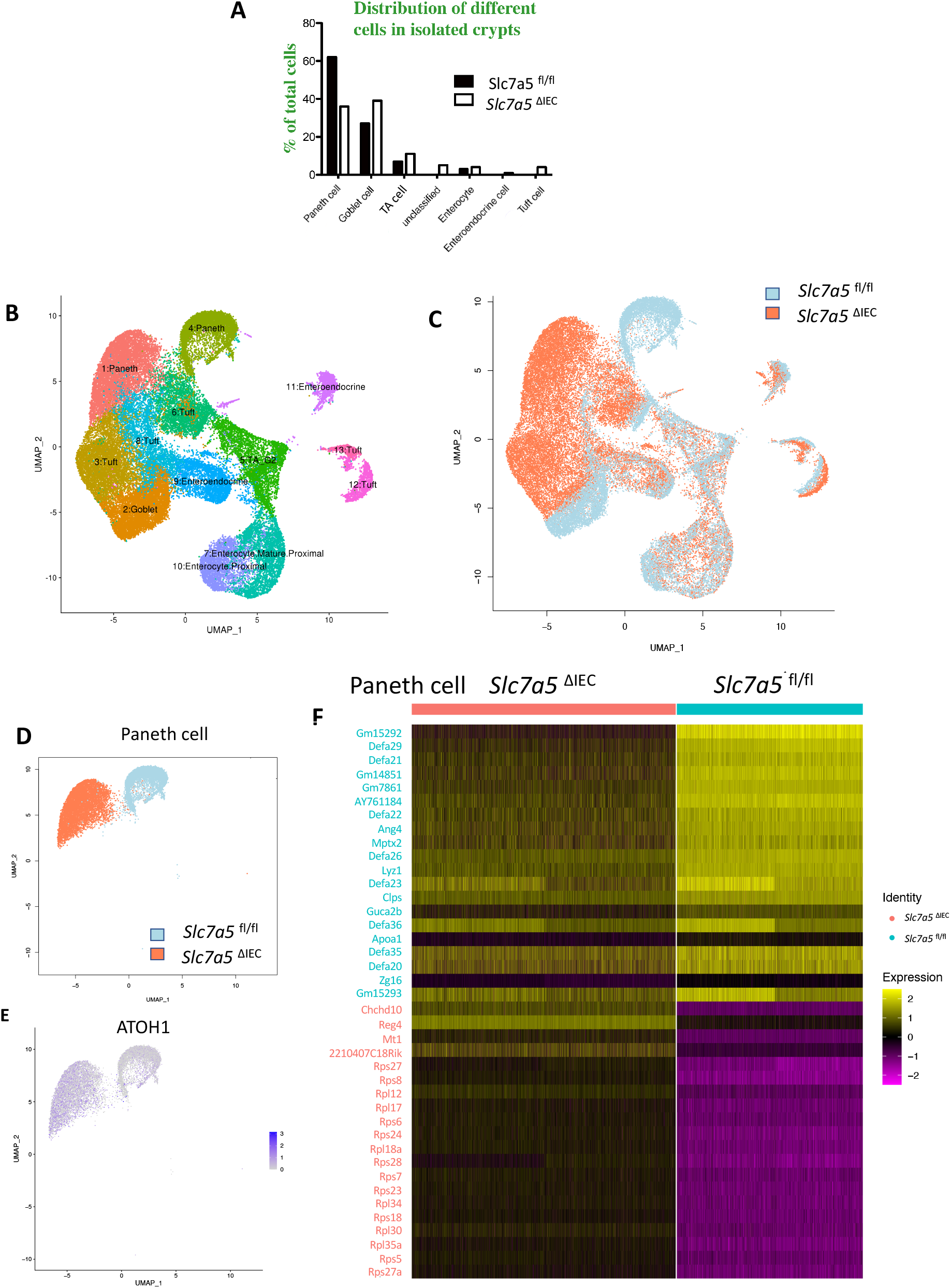
ScRNA-seq analysis of isolated small intestinal crypts reveals that *Slc7a5*^ΔIEC^ leads to expression of stem cell feature genes in Paneth cells. **A**. Percent distribution of small intestinal crypt cells among different cell types after from scRNA-seq. Epithelial cells from *Slc7a5*^fl/fl^ and *Slc7a5*^ΔIEC^ mice were clustered based on t-SNE plot of single cell RNA-seq data as shown Supplementary Fig. 2 and quantified as % of total cells sequenced for each animal type. **B**. Uniform manifold approximation and projection map (UMAP) showing different intestinal epithelial cell types (clusters) in different colors. All cells with scRNA-seq data from both *Slc7a5*^fl/fl^ and *Slc7a5*^ΔIEC^ mice were combined together and subjected to the analysis to show the distribution and location of different types of intestinal epithelial cells in different colors. **C**. The same as in B except the use of blue and orange colors for cells from *Slc7a5*^fl/fl^ and *Slc7a5*^ΔIEC^ mice, respectively. Note that most cells from the *Slc7a5*^fl/fl^ mice and *Slc7a5*^ΔIEC^ mice colocalized on the map. **D**. The view of the regions of the UMAP for Paneth cells from *Slc7a5*^fl/fl^ mice (blue dots) and *Slc7a5*^ΔIEC^ (orange dots) mice. Note that the distinct locations of the Paneth cells from *Slc7a5*^fl/fl^ mice and *Slc7a5*^ΔIEC^ mice suggests significant changes in gene expression patterns between them despite both having Paneth cell gene expression signatures. Similar views for other individual cell types (clusters) from *Slc7a5*^fl/fl^ mice and *Slc7a5*^ΔIEC^ mice are shown as Supplementary Fig. 3. **E**. The expression level of ATOH1 in Paneth cells as obtained from scRNA-seq was mapped on to the UMAP, showing higher levels in *Slc7a5*^ΔIEC^ mice than that in *Slc7a5*^fl/fl^ mice. F. Heatmap showing the relative expression levels of indicated genes in individual Paneth cells from *Slc7a5*^fl/fl^ and *Slc7a5*^ΔIEC^ mice. The genes shown here were features genes distinct for the Paneth cells from *Slc7a5*^fl/fl^ mice and *Slc7a5*^ΔIEC^ mice, with the top 20 for *Slc7a5*^fl/fl^ and bottom 20 for *Slc7a5*^ΔIEC^ Paneth cells.

To further analyze the scRNA-seq data, we combined all sequenced single cells from *Slc7a5*^fl/fl^ and *Slc7a5*^ΔIEC^ crypts and subjected them to unsupervised graph clustering (UMAP) with different epithelial cell types in different colors (Fig. 5B) or cells from *Slc7a5*^fl/fl^ and *Slc7a5*^ΔIEC^ mouse crypts in different colors (Fig. 5C). Comparison of Fig. 5B and 5C showed that most cell types from *Slc7a5*^fl/fl^ and *Slc7a5*^ΔIEC^ mouse crypts were co-localized. Indeed, when the locations of the cells of individual cell types on the UMAP were analyzed, we found that the cells from *Slc7a5*^fl/fl^ mice and *Slc7a5*^ΔIEC^ mice for most epithelial cell types were largely co-localized (Supplementary Fig. 3) with a small shift for the goblet cells and Tuft cells between the *Slc7a5*^fl/fl^ and *Slc7a5*^ΔIEC^ mice (Supplementary Fig. 3D, F). However, the locations of the Paneth cells from *Slc7a5*^fl/fl^ mice were distinct from the Paneth cells from *Slc7a5*^ΔIEC^ mice (Fig. 5D). When we compared the gene expression profiles of Paneth cells in *Slc7a5*^fl/fl^ and *Slc7a5*^ΔIEC^, we found 35 feature genes upregulated in *Slc7a5*^fl/fl^ Paneth cells compared to *Slc7a5*^ΔIEC^ Paneth cells and 843 feature genes upregulated in *Slc7a5*^ΔIEC^ Paneth cells compared to *Slc7a5*^fl/fl^ Paneth cells (Supplementary Data 1, 2), with the top 20 genes for each group visualized as a heatmap in Fig. 5F. Interestingly, among the top 20 genes expressed highly in *Slc7a5*^fl/fl^ mice were secretory granule genes, such as the *Defa* gene family, *Lyz* gene, and the *Gm* family gene, while those highly expressed in *Slc7a5*^ΔIEC^ mice contained many ribosomal genes, which are associated with regeneration and increased cell stemness (*30-33*). Finally, GO (Gene Ontology) analysis also showed that GO terms related to anti-microbial and immune responses were most significantly enriched among the genes expressed at higher levels in the *Slc7a5*^fl/fl^ Paneth cells than those in *Slc7a5*^ΔIEC^ Paneth cells, while GO terms related to metabolism and biosynthetic processes were most significantly enriched among the genes expressed at higher levels in the *Slc7a5*^ΔIEC^ Paneth cells than those in *Slc7a5*^fl/fl^ Paneth cells (Supplementary Fig 5 and Supplementary Data 3, 4). These again support the view that loss of SLC7A5 in intestinal epithelial cells causes dedifferentiation of the Paneth cells to gain stem cell features.

To test if the Paneth cells in the knockout mice indeed acquired or had increased stem cell features, we manually matched the 843 feature genes upregulated in *Slc7a5*^ΔIEC^ mice with Harber’s stem cell marker genes (*29*) and identified 5 genes shared between the two (Fig. S4). In addition, we analyzed all published stem cell marker genes known to facilitate regeneration of the intestinal epithelium (*34-41*), and found that ATOH1 was upregulated in *Slc7a5*^ΔIEC^ Paneth cells compared to the *Slc7a5*^fl/fl^ cells (Fig. 5E). Since ATOH1+ cells in crypts base are thought to be reserve stem cell, this observation also suggests that Paneth cells gained stem cell features, or had increased stemness, or de-differentiated in the *Slc7a5*^ΔIEC^ mice.

While scRNA-seq data showed a reduction in Paneth cells in *Slc7a5*^ΔIEC^ mice, the reduction was much less than what was observed from lysozyme-staining (Fig. 3A, B). This and the fact that Paneth cells in *Slc7a5*^ΔIEC^ mice appeared to have more stem cell features or be at least partially dedifferentiated prompted us to examine Paneth cells by electron microscopy. Under an electron microscope, the cells at the bottom of the crypts in *Slc7a5*^ΔIEC^ mice were not well organized and the number of secretory granules in Paneth cells were significantly decreased in comparison to *Slc7a5*^fl/fl^ mice (Fig. 6A). In addition, the rough endoplasmic reticulum (RER), where protein synthesis takes place, was also reduced in *Slc7a5*^ΔIEC^ Paneth cells (Fig. 6A), consistent with partial dedifferentiation of Paneth cells. Quantifications based on the ratio of nuclei to cytoplasm and the presence of granules in Paneth cells showed that the number of Paneth cells were similar between *Slc7a5*^fl/fl^ and *Slc7a5*^ΔIEC^ mice (Fig. 6B). These findings thus explained the apparent difference in the changes in Paneth cell number between scRNA-seq and lysozyme-staining since latter detects the presence of lysozyme in the granules. In addition, the electron microscopic analysis also showed that the crypt base stem cell numbers were similar between *Slc7a5*^ΔIEC^ mice and *Slc7a5*^fl/fl^ mice while mitotic cells at the crypt base were increased in *Slc7a5*^ΔIEC^ mice compared to *Slc7a5*^fl/fl^ mice (Fig. 6C, D), consistent with histological analyses above (Fig. 2C, Supplementary Fig. 1).

**Figure 6.**
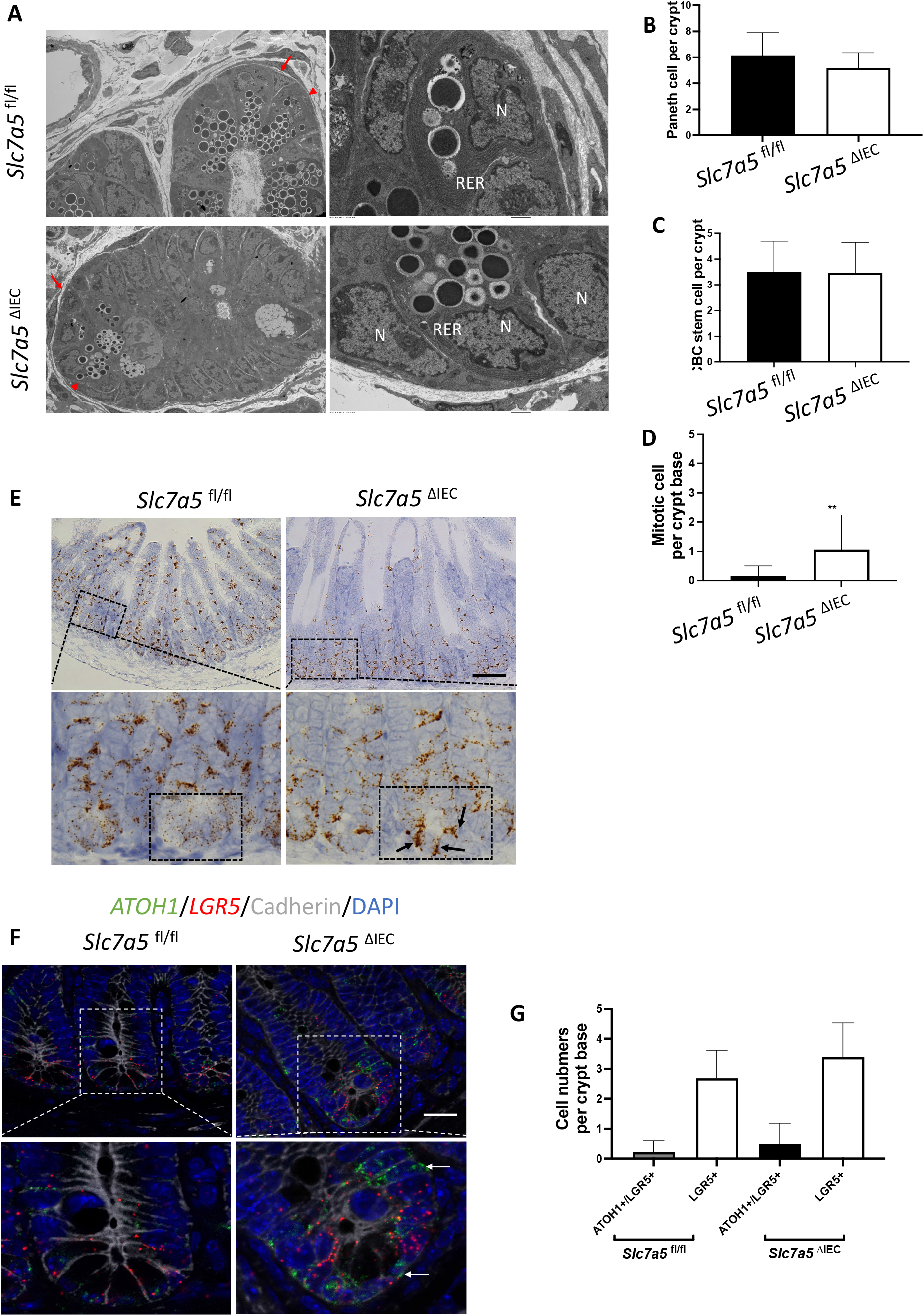
*Slc7a5*^ΔIEC^ secretory granules in Paneth cells and increases crypt base stem cell mitotic activity. **A**. Electronic microscopic images of Paneth cells in the small intestine of *Slc7a5*^fl/fl^ and *Slc7a5*^ΔIEC^ mice. Left panel: 1000x magnification. Note that crypt base in *Slc7a5*^ΔIEC^ mice was disorganized, had Paneth cells (red arrowhead) with reduced secretory granules, more mitotic cells. Right panel: 4000x magnification. Note that Paneth cells in *Slc7a5*^ΔIEC^ mice had reduced rough endoplasmic reticulum (right panel). N, nuclei; RER, rough endoplasmic reticulum. Red arrow, stem cells. **B**. Paneth cell numbers were similar between *Slc7a5*^fl/fl^ and *Slc7a5*^ΔIEC^ mice as quantified based on nucleus size and the present of secretory granules in Paneth cells on EM images. **C**. Crypt base stem cell numbers were similar between *Slc7a5*^fl/fl^ and *Slc7a5*^ΔIEC^ mice as quantified based on nucleus size and RER structure of cells on EM images. **D**. Crypt base mitotic cells increased in *Slc7a5*^ΔIEC^ mice as quantified on EM images. **E**. ATOH1 expression was higher in the crypt base of *Slc7a5*^ΔIEC^ mice compared to *Slc7a5*^fl/fl^ mice as detected by single molecule in situ hybridization. The region with a dashed box in each lower panel represents a crypt base, showing increased ATOH1 expression in the *Slc7a5*^ΔIEC^ mice. Scale bars, 100µm. **F**. Co-expression of ATOH1 and LGR5 in some crypt base stem cells in *Slc7a5*^fl/fl^ and *Slc7a5*^ΔIEC^ mice. Single molecule in situ hybridization were performed with a mixture of probes for ATOH1 (green), LGR5 (red), and cadherin (gray) on intestinal cross-sections and the DNA was stained blue. The boxed regions in the top panel were enlarged in the bottom panel. Scale bars, 50µm. White arrow, ATOH1+ cells. **G**. LGR5 and ATOH1 were not co-expressed in most cells. A cell was considered to be positive for an mRNA if at least 5 dots (single molecular hybridization signal) for the mRNA was present in a cell. Results were shown as mean ± SD.

To further investigate the potential dedifferentiation of the Paneth cells in *Slc7a5*^ΔIEC^ mice, we used single molecule *in situ* hybridization (smISH) and immunofluorescence to detect ATOH1, LGR5, Ki67 and lysozyme at crypt base. The smISH analysis revealed that ATOH1, whose high-level expression has been implicated to a marker for reserved stem cells (*35, 36*), was highly expressed in the crypt base of *Slc7a5*^ΔIEC^ mice compared to *Slc7a5*^fl/fl^ mice (Fig. 6E). By co-staining for LGR5, a marker for CBC stem cells, or lysozyme, a marker for mature Paneth cells, we found that ATOH1 mRNA was mostly co-expressed with lysozyme (Supplementary Fig. 6A, D) (note that the fraction of ATOH1-positive cells with lysozyme was decreased a little in *Slc7a5*^ΔIEC^ mice, likely due to reduced granules that made some Paneth cells not positive for lysozyme detection in *Slc7a5*^ΔIEC^ mice), but not with LGR5 (Fig. 6F, G). Thus, ATOH1 is largely a marker for Paneth cells at crypt base but its expression was increased in the Paneth cells in *Slc7a5*^ΔIEC^ mice, in agreement with the scRNA-seq data (Fig. 5E). In addition, co-staining for Ki67 showed that in *Slc7a5*^ΔIEC^ mice, there were proliferating cells at the crypt base with ATOH1 expression (Supplementary Fig. 6B, E), likely due to increased proliferation of some stem cells expressing both LGR5 and ATOH1 (Fig. 6F, G). Furthermore, 80% LGR5-positive cells were stained positively for Ki67, compared to 30% in *Slc7a5*^fl/fl^ mice (Supplementary Fig. 3), indicating that *Slc7a5*^ΔIEC^ increased the proliferation of CBC stem cells, consistent with the data from Ki67 staining alone (Fig. 2C).

### Loss of SLC7A5 in intestinal epithelium sensitizes mice to DSS-induced colitis

It is well known that Paneth cells and goblet cells play critical roles maintaining intestinal integrity and defense. Paneth cells secrete AMPs to protect the epithelium from pathogens while the goblet cells secret mucus to block the access of the microbiome to mucosa and to allow the accumulation of high concentration of AMPs. The reduction of mature secretory cells, suggests that the mutant mice may have altered inflammatory response and are more susceptible to IBD or colitis. To test this possibility, we compared the expression of IBD-related inflammatory gene mRNAs between *Slc7a5*^fl/fl^ mice and *Slc7a5*^ΔIEC^ mice by using the normalized total read counts from scRNA-seq data and found that the expression of many such genes were increased in the intestinal crypt of *Slc7a5*^ΔIEC^ mice compared to that in *Slc7a5*^fl/fl^ mice (Supplementary Fig. 7), further suggesting that *Slc7a5*^ΔIEC^ mice may be more susceptible to inflammatory diseases in the intestine. To test this possibility, we subjected *Slc7a5*^ΔIEC^ and *Slc7a5*^fl/fl^ mice to dextran sodium sulphate (DSS) treatment to induced intestinal inflammation (Fig. 7A). After the treatment, the colonic epithelium of *Slc7a5*^ΔIEC^ mice had much more severe necrotic disruption/loss (Fig. 7B), accompanied with thicker submucosal layers, compared to the *Slc7a5*^fl/fl^ mice. Consistent with the damage to the intestine, the *Slc7a5*^ΔIEC^ mice lost weight faster throughout the treatment (Fig. 7C). Using a well-established histology scoring method for colitis (*42*), we found that the loss of SLC7A5 in the intestinal epithelium led to much more severe DSS-induced colitis compared to *Slc7a5*^fl/fl^ mice. Thus, *Slc7a5*^ΔIEC^ renders mice more susceptible to experimentally induced colitis.

**Figure 7.**
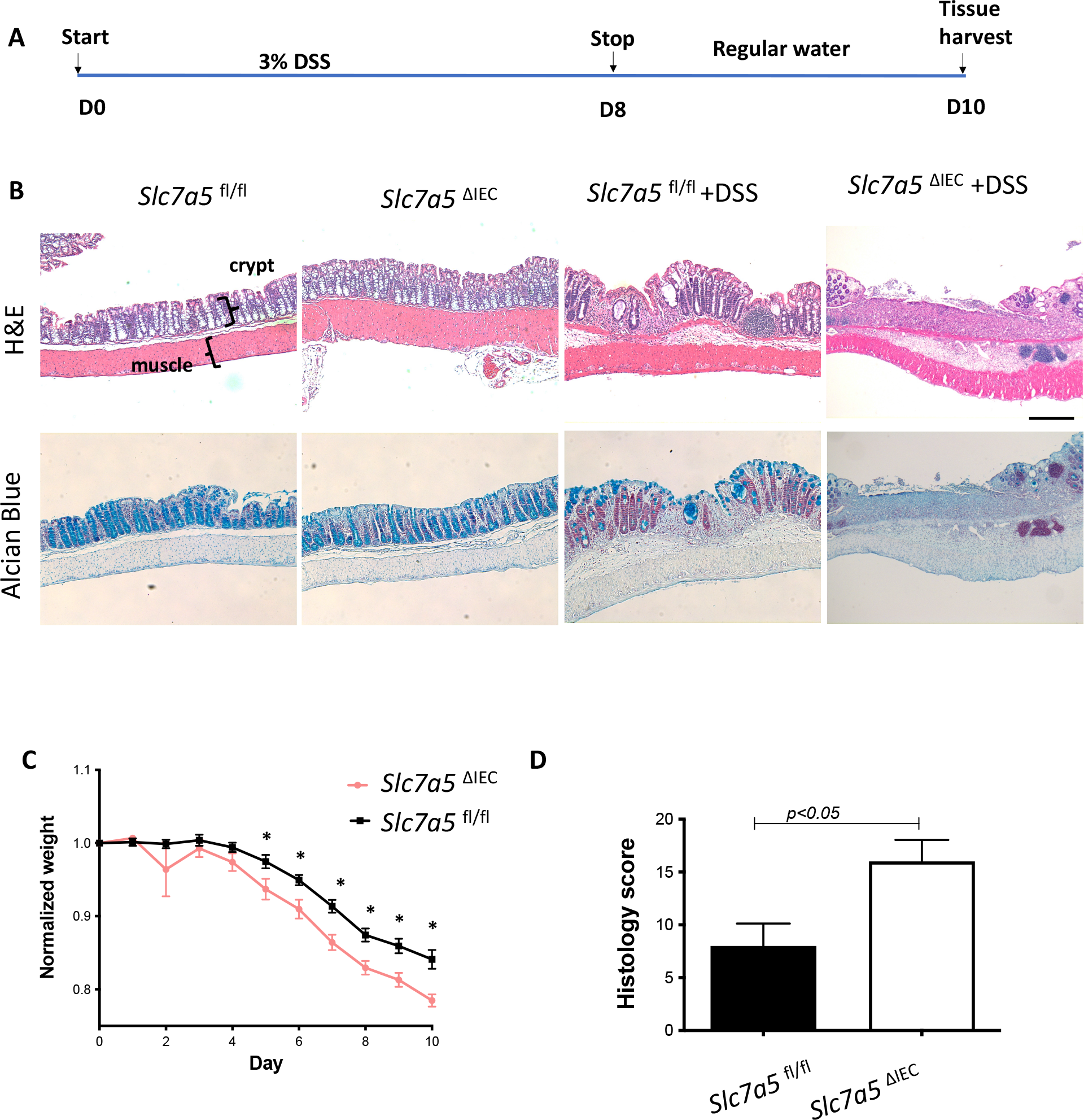
*Slc7a5*^ΔIEC^ mice are more susceptible to DSS-induced colitis. **A**. Schematic diagram for the 10 days DSS-induced colitis experiment. **B**. HE and Alcian blue staining showing colon structures for *Slc7a5*^fl/fl^ and *Slc7a5*^ΔIEC^ mice with or without DSS treatment at the end of the experiment. Note the more severe disruption or loss of the epithelial structure in *Slc7a5*^ΔIEC^ mice. Scale bars, 100µm. **C**. Weight loss was more severe in *Slc7a5*^ΔIEC^ mice during DSS treatment (n=10). * p<0.05. D. *Slc7a5*^ΔIEC^ mice had more severe colitis as judged based on histology sores. Results were shown as mean ± SD

## Discussion

Amino acids are key inputs of mTORC1 signaling and thus many studies on mTORC1 function in intestinal homeostasis have used calorie restriction as the model. However, results from different studies are sometimes inconsistent. Notably, Yousefi et al. (*20*) showed that calorie restriction enhanced intestinal epithelial regenerative capacity through a cell-autonomous mechanism by directly downregulating mTORC1 in reserve stem cell under basal conditions but allowing for robust mTORC1 activation in response to radiation injury to a greater extent than in *ad libitum*-fed mice. However, Igarashi et al. (*18*) and Yilmaz et al. (*19*) showed that caloric restriction mainly increased CBC stem cell pool through a non-cell-autonomous mechanism by directly downregulating mTORC1 in Paneth cells, which secrete cyclic ADP ribose to activate neighbor LGR5-positive stem cells, causing their expansion. Despite the difference, both studies suggest nutrition shortage can affect different cell types via mTORC1 signaling. In addition, it was reported that disruption of mTORC1 signaling caused a reduction in Paneth cell number and led to less regeneration capacity after radiation injury (*22*). These various studies suggest that alteration of mTORC1 signaling can influence epithelial cell regeneration.

The present study provides further evidence on the role of mTORC1 signaling in intestinal homeostasis by introducing a genetic model in which mTORC1 signaling is decreased via knockout of the amino acid transporter SLC7A5 in intestinal epithelial cells. We showed that loss of SLC7A5 caused Paneth cell de-differentiation, leading to disruption in secretory function through a reduction in secretory granules. The overall phenotype of *Slc7a5*^ΔIEC^ mice is similar to that of the mTORC1 intestinal epithelial cell-specific knockout mouse (*22*).

The mTORC1 signaling is generally associated with cell proliferation. As an activator of mTORC1 signaling through amino acid transport, SLC7A5 may be expected to be associated with cell proliferation. For example, cancer cells require a massive nutrient supply and SLC7A5, as an amino acid transporter, has been widely studied in various cancers, highlighting the importance of amino acid transport in cell proliferation and tumor growth (*43-45*). In addition, in KRAS-mutant colorectal cancer, *Slc7a5* disruption abrogates tumor cell growth (*46*). Our *Slc7a5*^ΔIEC^ mice expectedly had a reduction in mTORC1 signaling in the intestinal crypts but surprisingly led to increased proliferation of crypt base stem cells and transit amplifying cells. It is puzzling how the decrease of mTORC1 causes an overall increase in cell proliferation. Our scRNA-seq data showed that Paneth cells had increased expression of stem cell feature genes, like ATOH1 and ribosomal genes. Ribosomal genes have been widely linked to carcinogenesis and stem cells (*30, 32, 33*). Paneth cells have been shown to acquire a stem cell-like transcriptome post-injury and are able to regenerate the whole intestinal epithelium through activation of Notch signaling (*47*). ATOH1 functions as an essential downstream marker of Notch pathway (*48, 49*) and ATOH1+ cells can regenerate the whole intestinal epithelium after injury based on a physical lineage tracing study (*36*). Thus, it is possible that loss of intestinal epithelial cell SLC7A5 causes Paneth cell dedifferentiation that may eventually lead to highly proliferative stem cells and transit amplifying cells, although we did not observe a significant increase in crypt base stem cells.

Alternatively, Paneth cells could influence cell proliferation indirectly. It is well known that Paneth cells can function as stem cell niche to influence stem cell identity and proliferation (*18, 19, 50, 51*). Thus, the dedifferentiation of Paneth cells due to *Slc7a5*^ΔIEC^ may alter the niche and/or other secreted factors to increase the proliferation of stem cells and transit amplifying cells (Fig. 8). It is also possible that increased cell proliferation in *Slc7a5*^ΔIEC^ mice may be secondary to epithelial damage caused by Paneth cell dysfunction (Fig. 8). As have shown herein, *Slc7a5*^ΔIEC^ mice are prone to epithelial damage, such as DSS-induced colitis, due to reduced Paneth cell and goblet cell function. Such loss of protection from AMPs and other external factors (*8-10, 52*) may cause intestinal inflammatory stress, which in turn leads to increased epithelial cell death. The resulting loss of epithelial cells can then generate a compensatory increase in cell proliferation to maintain intestinal homeostasis (Fig. 8) (*53-55*). Such a mechanism is consistent with observation from an enteritis model due to knockout XBP1 (*56*). XBP1 is an endoplasmic reticulum (ER) stress regulator. Knockout XBP1 can lead to enteritis, which causes Paneth cell dysfunction and increased cell proliferation in order to repair the epithelial damage due to enteritis. Both mTORC1 and ER stress have been genetically linked (*57*) and *Xbp1*^-/-^ mice share a similar phenotypes with the *Slc7a5*^ΔIEC^ mice, suggesting possible common involvement of Paneth cell dysfunction in influencing cell proliferation. Clearly, further studies are needed to decipher how SLC7A5 regulates intestinal homeostasis through mTORC1 signaling. Regardless of the exact mechanism, our findings highlight an important role of amino acid transport and/or nutritional status in regulated Paneth cell function to affect intestinal homeostasis and pathogenesis.

**Figure 8.**
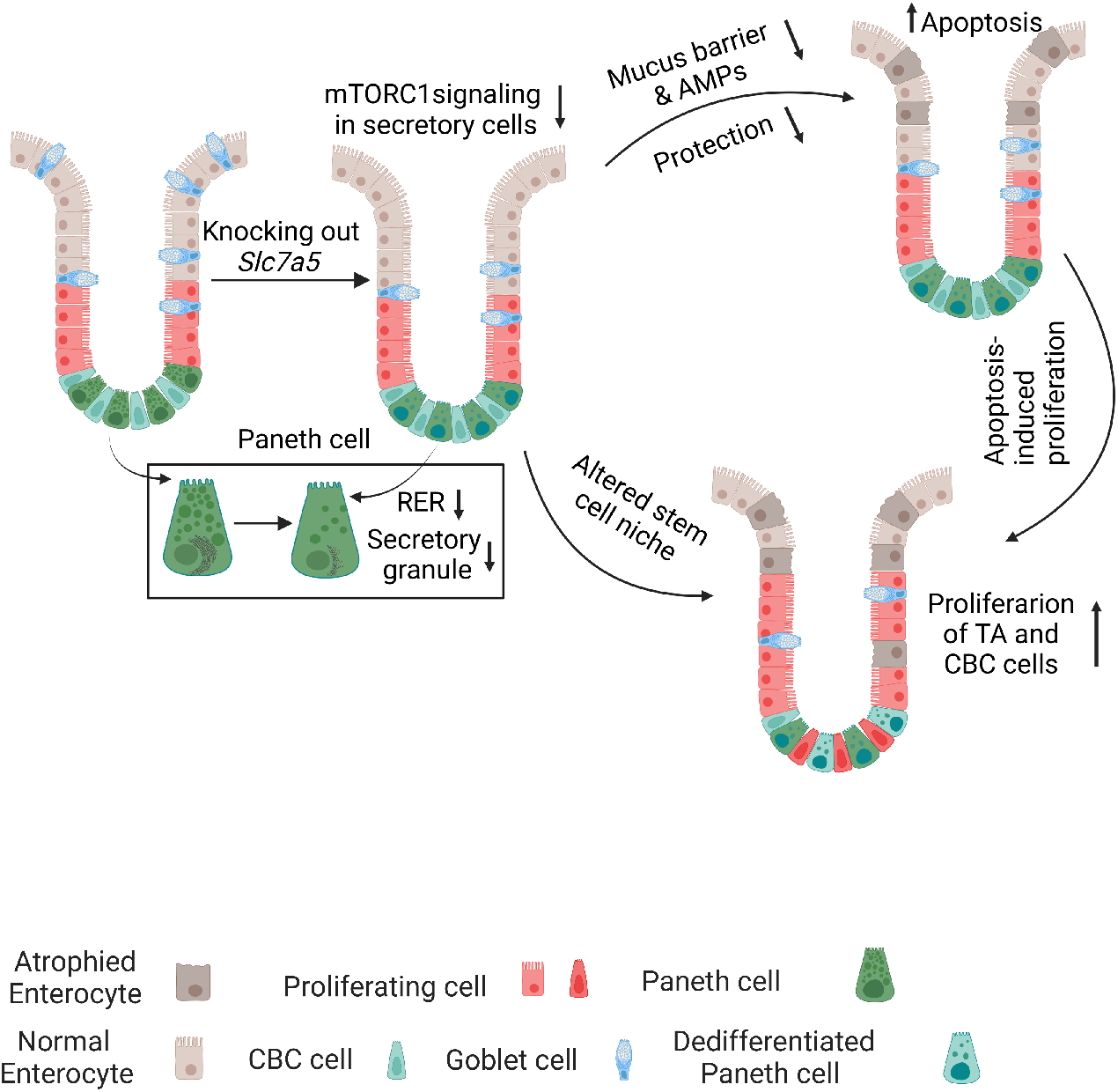
A model for the effect of *Slc7a5*^ΔIEC^ on intestinal homeostasis and function. Intestinal epithelial specific knockout of SLC7A5 causes a reduction of mTORC1 signaling mainly in the secretory cell lineage, particular Paneth cells in the crypt base, which have reduced RER and secretory granules. This leads to a loss in secretory cell functions, such as reduced mucus barrier and secretion of anti-microbial peptides (AMPs). This in turn weakens protection against pathogens and/or mechanical stresses from food. The consequences are to increase intestinal inflammatory stress that can result in increased damage due to external challenges like DSS treatment, and/or and increased epithelial apoptosis, which can result in proliferation of TA and/or CBC stem cells to maintain epithelial homeostasis. In addition, Paneth cell dedifferentiation in the knockout mice, as reflected by the reduction in secretory granules and gains in stem cell feature gene expression, will likely alter stem cell niche and thus contribute to increased cell proliferation in the crypt.

## Materials and Methods

### Animals

*Slc7a5*^fl/fl^ mice were previously described (*16*). Villin-Cre (The Jackson Laboratory; stock no. 021504) mice, backcrossed to C57BL/6J mice for more than 10 generations, were bred with *Slc7a5*^fl/fl^ mice to produce *Slc7a5*^ΔIEC^ mice (Fig. 1A). *Slc7a5*^fl/fl^ mice were use as the wild type controls for *Slc7a5*^ΔIEC^ mice. The *Slc7a5*^ΔIEC^ and *Slc7a5*^fl/fl^ mice were sex- and age-matched littermates and cohoused. All mice were maintained in accordance with the NIH animal facility guidelines for laboratory animal research. All animal care and treatments were done as approved by Animal Use and Care Committee of *Eunice Kennedy Shriver* National Institute of Child Health and Human Development, National Institutes of Health.

### Measurements for small intestine, intestinal crypts, villi, and body

The body length and body weight were measured on euthanized *Slc7a5*^fl/fl^ and *Slc7a5*^ΔIEC^ mice from the tip of the mouth to the base of the tail. The length of dissected small intestine was measured from the beginning of the duodenum to the end of the ileum. The length of the villus was measured from the mouth of the crypt to the tip of the villus on H&E-stained intestinal cross-sections. For the isolation of intestinal epithelial cells, the intestine was cut into 2-5 mm segments and incubated in phosphate-buffered saline (PBS) containing 20 mM EDTA for 30 min on ice on a rocker. Epithelial cells were released by vigorous shaking or pipetting and passed through a 70 mm strainer to collect enriched crypts or all the filtrate was used to collect intestinal epithelial cells, which were then washed with cold PBS containing 0.1% bovine serum albumin (BSA).

### Mouse intestinal organoids

IntestiCult Organoid Growth Medium (mouse) (Stem Cell Technologies) was used to establish and culture mouse intestinal organoids. Briefly, the mice were killed and the whole small intestine was harvested from each mouse. The intestine was flushed gently with cold PBS. Scissors were used to make a longitudinal incision along the entire length of the intestine, and the intestine was then cut into 2-5 mm pieces. A 20-ml serological pipette was used to wash the intestinal pieces by pipetting up and down three times. The intestinal pieces were sedimented by centrifugation and re-suspended into fresh PBS. This wash procedure was repeated 15-20 times. Then, the intestinal pieces were suspended in 20 mM EDTA for 30 min on ice on a rocker before pelleting by centrifugation. They were then resuspended in 20 ml of cold PBS containing 0.1% BSA and pipetted up and down three times. The supernatant was then passed through a 70-mm strainer to collect the enriched crypts in the filtrate. This procedure was repeated 3-6 times to obtain enough crypts for organoid culture. The quality of the crypts in the suspension was assessed, and the number of crypts were counted with an inverted microscope. Crypts were then cultured in domes made with a mixture of Matrigel (Corning) and IntestiCult Organoid Growth Medium (1:1). Fifty microliter of the mixture was pipetted into a 24-well plate. Next, the plate was incubated at 37 °C for 10 min to allow the Matrigel to solidify. Finally, 500µl complete IntestiCult Organoid Growth Medium were added to each well and the plate was then incubated at 37 °C and 5% CO_2_ for 7 days with the medium changed every other day. For immunoblotting assays, 10 mM BCH were incubated for three days before harvest; 20 µM rapamycin, or 10 µM MHY1485 for overnight.

### EdU staining

Proliferating cells were labeled with 5-ethynyl-2’-deoxyuridine (EdU)-incorporation. Click-iT^™^ EdU Cell Proliferation Kit for Imaging (Thermo Fisher Scientific, Alexa Fluor^™^ 594 dye) was applied following manufacturer’s protocol. Briefly, for cell proliferation assay, mice were intraperitoneally injected with EdU (1 mg per mouse) two hours before killing. In EdU pulse-chase assay, mice were intraperitoneally injected with EdU (1 mg per mouse), and the mice were killed at 2 h, 8 h, 24 h, 48 h or 72 h, after injection. The EdU-labeled intestine was isolated for paraffin sections (5 µm) as described above. The intestinal paraffin sections were baked at 60 °C for 1 h followed by deparaffinized with xylene and rehydrated through a graded series of ethanol. Then the sections were incubated with Click-iT^®^ Plus reaction cocktail for 30 minutes at room temperature, washed once with 1 × PBST (1 × PBS and 0.05% Tween-20), mounted on glass slides with ProLong^®^ Gold Antifade Mountant with DAPI (Thermo Fisher Scientific). The fluorescent pictures for different colors and different sections were taken under the same settings and then analyzed by using ImageJ at the same setting to count the positive cells.

### Immunohistochemistry

The intestine was removed from age-matched *Slc7a5*^fl/fl^ and *Slc7a5*^ΔIEC^ littermates, flushed with ice-cold 1xPBS buffer, and fixed in 4% formaldehyde (and if needed, stored at 4 °C), followed by embedding in paraffin and then cutting to 5 µm sections. For H&E staining, the 5 µm sections were stained with hematoxylin and eosin following manufacturer’s protocol (Sigma) and analyzed under a bright field microscope. For immunofluorescent staining, four to five 5 μm paraffin sections/animal were deparaffinized in xylene and rehydrated in a series of different concentrations of ethanol. Antigen retrieval was performed by boiling in an antigen retrieval buffer (1 mM Tris, 1 mM EDTA and 0.05% Tween-20) for 3 min at 125 °C followed by washing the slides under running water and rinsing them in 1xTBS-Tween (Tris buffered saline plus 0.1% tween-20) for 5 min. After incubation in blocking buffer (10% normal goat serum in PBS) for 1 h at room temperature, the primary antibody was added and the slides were incubated at 4 °C overnight. The slides were then washed in 1xTBS-Tween and subsequently incubated with a fluorescence-labeled secondary antibody for 1 h at room temperature, before the slides were washed three times with 1x TBS-Tween and covered with DAPI-containing mounting medium to counterstain the DNA. The fluorescent pictures for different colors and/different sections were taken under the same settings. The fluorescent pictures were analyzed by using the Image J at the same setting to count positive cell numbers. The primary antibodies used were anti-lysozyme antibody (Abcam, and Dako) and Ki67 (Cell Signaling Technology). For immunohistochemical staining, paraffin sections of the intestine were incubated with the primary antibody for 1 h at room temperature, followed by washing steps with PBS as described above for immunohistochemistry. The HRP/DAB (ABC) Detection kit (Abcam) was used to detect the signals by following the manufacturer’s protocol. The brown Paneth cells in the crypt were then visually counted double-blindly.

### Apoptosis detection

Apoptosis was analyzed with terminal deoxynuclotidyltransferase-mediated dUTP nick-end labeling (TUNEL; Roche) following the manufacturer’s protocol. Briefly, the TUNEL *in situ* Cell Death Detection Kit was used on 5 μm paraffin sections of the intestine. Sections were dewaxed with xylene and rehydrated with a series of different concentrations of ethanol. Antigen retrieval was performed by microwaving the sections (700W; 1.5 min) in sodium citrate buffer (pH 6.0) followed by rinsing in PBS. The sections were incubated for 30 min in 0.1 M Tris-HCL pH 7.5, containing 1.5% BSA, and 20% normal bovine serum, washed in PBS, and incubated with TUNEL reaction mixture at 37 °C for 1 h. After removing the TUNEL reaction mixture, sections were washed in PBS several times and counterstained with DAPI. The fluorescent pictures for different colors and/different sections were taken under the same settings and then analyzed by using Image J at the same setting to count the positive cell numbers.

### Alcian blue staining for goblet cells

Paraffin sections of the intestine were stained with Alcian Blue Kit (Abcam) following the manufacturer’s protocol. The blue goblet cells in the crypt and villus were counted visually or analyzed by using Fiji Image J.

### In situ hybridization

*In situ* hybridization with RNAscope 2.5 HD Reagent Kit-Brown (322371; Advanced Cell Diagnostics, Newark, CA) or RNAscope Multiplex Fluorescent V2 Assay (323100; Advanced Cell Diagnostics, Newark, CA) was performed on 5 μm, formalin-fixed, paraffin-embedded sections according to manufacturer’s instructions. The RNAscope probes used were LGR5, OLFM4, ATOH1, the negative control probe *DapB*, and the positive control probe *Ppib*. The positive signals in crypt were quantified by using Fiji Image J or Imaris.

### SS-induced colitis

*Slc7a5*^ΔIEC^ and *Slc7a5*^fl/fl^ mice were treated with 3% DSS in water for 8 days and changed to regular water for 2 days. The weights of the mice were monitored at the same time point every day. The mice were killed on indicated days, and the colon was fixed for histology assessment of the severity of the colitis. The range along the intestine and depth of epithelium damage, mononuclear cell infiltration, loss of goblet cells was used for scoring the severity of the colitis based on published scoring criteria (*42*).

### Western blot

Total proteins from intestinal epithelial cells or intestinal organoids were extracted with M-PER Mammalian Protein Extraction Reagent (Thermo Fisher Scientific) with proteinase inhibitor (Roche) and phosphatase inhibitor (Thermo Fisher Scientific) and then subjected to SDS-polyacrylamide gel electrophoresis, followed by transferring to polyvinylidene difluoride membranes and subsequent immunoblotting assay. Quantification was performed with Li-COR analysis system (LI-COR Bioscience). The antibodies used were anti-phospho-p70S6K1 (Cell Signaling Technology), anti-p70S6K1 (Cell Signaling Technology), anti-phospho-S6 (Cell Signaling Technology), anti-S6 (Cell Signaling Technology), anti-β-actin (Abcam).

### scRNA-seq and data analysis

The enriched intestinal crypts as isolated above were washed twice in PBS, centrifuged at 300 g for 3 min, and the epithelial cells dissociated with TrypLE express (Invitrogen) for 1 min at 37 °C. The dissociated single cells were stained with an EpCAM (Thermo Fisher Scientific), CD45 (Thermo Fisher Scientific), and propidium iodide (Thermo Fisher Scientific). The EpCAM+ CD45-live cells were sorted by flow cytometry and cells was counted. The estimated ideal number of cells were used for scRNA-seq according to manufacturer’s instructions (10x Genomics kit, chromium single cell 3’ reagent kits V2). Single cell sequencing was performed with Illumina HiSeq2500 sequencer at Center for Cancer Research Sequencing Facility, National Cancer Institute, National Institutes of Health. For analysis of scRNA-seq data, a total of 25,776 cells, 6559 cells from *Slc7a5*^fl/fl^ mice and 19217 cells from *Slc7a5*^ΔIEC^ mice, after filtering and removing the immune clusters, were used. 10x Genomics Cell Ranger software (version 3.1.0) was used for single cell demultiplexing, alignment, filtering, barcode counting, and UMI counting. Then the counting outputs were processed with Seurat package (v3.2.3) under R (version 4.0.3) following the standard workflow (https://satijalab.org/seurat/articles/pbmc3k_tutorial.html). Cell type identity was assigned to cell clusters based on feature genes of intestinal epithelial cells subtypes(*29*). To compare intestinal epithelial cell subtypes between *Slc7a5*^ΔIEC^ and *Slc7a5*^fl/fl^ mice, the counting data of both *Slc7a5*^ΔIEC^ and *Slc7a5*^fl/fl^ mouse samples were aggregated as one with 10x Genomics Cell Ranger software and processed with R Seurat package and the same workflow as above. After assigning cell types to clustered cells, differentially-expressed genes between different clusters were identified with FindMarkers function in R Seurat package with setting: only.pos=True, min.pct=0.25, and logfc.threshold=0.25.

### Electronic microscopy

Mice were transcardially perfused fixed with 4% PFA with 2.5% glutaraldehyde, made in PBS buffer, pH 7.4. Small intestines were removed and left to post-fix overnight in the same fixative at 4 °C. All tissue samples were then rinsed in 0.1M sodium cacodylate buffer. The following processing steps were carried out using the variable wattage Pelco BioWave Pro microwave oven (Ted Pella, Inc., Redding, CA.): post-fixed in 1% osmium tetroxide made in 0.1 M sodium cacodylate buffer, rinsed in double distilled water (DDW), 2% (aq.) uranyl acetate enhancement, DDW rinse, ethanol dehydration series up to 100% ethanol and propylene oxide, followed by a Embed-812 resin (Electron Microscopy Sciences, Hatfield, PA.) infiltration series up to 100% resin. The epoxy resin was polymerized for 20 h in an oven set at 60 °C. Ultra-thin sections were cut on a Leica EM-UC7 Ultramicrotome (90 nm). Thin sections were picked up and placed on 200 mesh cooper grids and post-stained with uranyl acetate and lead citrate. Imaging was accomplished using a JEOL-1400 Transmission Electron Microscope operating at 80kV and an AMT BioSprint-29 camera.

### Statistical analysis

Representative images of at least two independent experiments with each having at least three mice for each sample group were shown. All statistical analyses and graphs were performed/generated by using GraphPad Prism version 8.0 (GraphPad Software, La Jolla, CA). Student’s t test was used to examine the differences between groups and a P < 0.05 was considered statistically significant. In the analysis of histology, multiple sections from each animal were used and 10 to 45 sections in total were analyzed for each group. All data were expressed as mean ± SD.

## Supporting information

supplementary image1-7

## Acknowledgments

Electron Microscopy was performed at the NICHD Microscopy Imaging Core with the assistance of Dr. Louis Dye.

Figure 8 is created with BioRender.com.

## Author contributions

Conceptualization: LB, YBS Methodology: LB, LF, HZ, ZC, LS, ZP Visualization: LB, YS, HZ Supervision: YBS, BS, CW, FG Writing—original draft: LB, YBS Writing—review editing: all authors

## Data and materials availability

All data are available in the main text or the supplementary materials. ScRNA-seq data have been deposited in Gene Expression Omnibus database with the accession code GSE216930.

## References

1. J. Beumer, H. Clevers, Cell fate specification and differentiation in the adult mammalian intestine. Nat Rev Mol Cell Biol 22, 39–53 (2021).

2. L. W. Peterson, D. Artis, Intestinal epithelial cells: regulators of barrier function and immune homeostasis. Nat Rev Immunol 14, 141–153 (2014).

3. J. Gao, K. Xu, H. Liu, G. Liu, M. Bai, C. Peng, T. Li, Y. Yin, Impact of the Gut Microbiota on Intestinal Immunity Mediated by Tryptophan Metabolism. Front Cell Infect Microbiol 8, 13 (2018).

4. K. J. Maloy, F. Powrie, Intestinal homeostasis and its breakdown in inflammatory bowel disease. Nature 474, 298–306 (2011).

5. A. J. M. Santos, Y. H. Lo, A. T. Mah, C. J. Kuo, The Intestinal Stem Cell Niche: Homeostasis and Adaptations. Trends Cell Biol 28, 1062–1078 (2018).

6. N. Barker, S. Bartfeld, H. Clevers, Tissue-resident adult stem cell populations of rapidly self-renewing organs. Cell Stem Cell 7, 656–670 (2010).

7. H. Gehart, H. Clevers, Tales from the crypt: new insights into intestinal stem cells. Nat Rev Gastroenterol Hepatol 16, 19–34 (2019).

8. M. E. Johansson, H. Sjövall, G. C. Hansson, The gastrointestinal mucus system in health and disease. Nature reviews Gastroenterology hepatology 10, 352–361 (2013).

9. J. Shi, Defensins and Paneth cells in inflammatory bowel disease. Inflammatory bowel diseases 13, 1284–1292 (2007).

10. Y. Nakanishi, M. Reina-Campos, N. Nakanishi, V. Llado, L. Elmen, S. Peterson, A. Campos, S. K. De, M. Leitges, H. Ikeuchi, M. Pellecchia, R. S. Blumberg, M. T. Diaz-Meco, J. Moscat, Control of Paneth Cell Fate, Intestinal Inflammation, and Tumorigenesis by PKClambda/iota. Cell Rep 16, 3297–3310 (2016).

11. N. Singh, G. F. Ecker, Insights into the Structure, Function, and Ligand Discovery of the Large Neutral Amino Acid Transporter 1, LAT1. Int J Mol Sci 19, p(2018).

12. M. Scalise, M. Galluccio, L. Console, L. Pochini, C. Indiveri, The Human SLC7A5 (LAT1): The Intriguing Histidine/Large Neutral Amino Acid Transporter and Its Relevance to Human Health. Front Chem 6, 243 (2018).

13. J. W. A. R. a. P. M. Taylor, Role of the System L permease LAT1 in amino acid and iodothyronine transport in placenta. Biochemical Society 725, 719–725 (2002).

14. K. Ozaki, T. Yamada, T. Horie, A. Ishizaki, M. Hiraiwa, T. Iezaki, G. Park, K. Fukasawa, H. Kamada, K. Tokumura, The L-type amino acid transporter LAT1 inhibits osteoclastogenesis and maintains bone homeostasis through the mTORC1 pathway. Science signaling 12, eaaw3921 (2019).

15. L. V. Sinclair, J. Rolf, E. Emslie, Y. B. Shi, P. M. Taylor, D. A. Cantrell, Control of amino-acid transport by antigen receptors coordinates the metabolic reprogramming essential for T cell differentiation. Nat Immunol 14, 500–508 (2013).

16. N. Poncet, F. E. Mitchell, A. F. Ibrahim, V. A. McGuire, G. English, J. S. Arthur, Y. B. Shi, P. M. Taylor, The catalytic subunit of the system L1 amino acid transporter (slc7a5) facilitates nutrient signalling in mouse skeletal muscle. PLoS One 9, e89547 (2014).

17. T. E. Harris, M. O. Thorner, Caloric restriction in mTORC1 control of intestinal homeostasis. Cell Metab 16, 6–8 (2012).

18. M. Igarashi, L. Guarente, mTORC1 and SIRT1 Cooperate to Foster Expansion of Gut Adult Stem Cells during Calorie Restriction. Cell 166, 436–450 (2016).

19. O. H. Yilmaz, P. Katajisto, D. W. Lamming, Y. Gultekin, K. E. Bauer-Rowe, S. Sengupta, K. Birsoy, A. Dursun, V. O. Yilmaz, M. Selig, G. P. Nielsen, M. Mino-Kenudson, L. R. Zukerberg, A. K. Bhan, V. Deshpande, D. M. Sabatini, mTORC1 in the Paneth cell niche couples intestinal stem-cell function to calorie intake. Nature 486, 490–495 (2012).

20. M. Yousefi, A. Nakauka-Ddamba, C. T. Berry, N. Li, J. Schoenberger, K. P. Simeonov, R. J. Cedeno, Z. Yu, C. J. Lengner, Calorie Restriction Governs Intestinal Epithelial Regeneration through Cell-Autonomous Regulation of mTORC1 in Reserve Stem Cells. Stem Cell Reports 10, 703–711 (2018).

21. N. Poncet, P. A. Halley, C. Lipina, M. Gierlinski, A. Dady, G. A. Singer, M. Febrer, Y. B. Shi, T. P. Yamaguchi, P. M. Taylor, K. G. Storey, Wnt regulates amino acid transporter Slc7a5 and so constrains the integrated stress response in mouse embryos. EMBO Rep 21, e48469 (2020).

22. L. L. Sampson, A. K. Davis, M. W. Grogg, Y. Zheng, mTOR disruption causes intestinal epithelial cell defects and intestinal atrophy postinjury in mice. FASEB J 30, 1263–1275 (2016).

23. M. K. Holz, B. A. Ballif, S. P. Gygi, J. Blenis, mTOR and S6K1 mediate assembly of the translation preinitiation complex through dynamic protein interchange and ordered phosphorylation events. Cell 123, 569–580 (2005).

24. M. Zhang, F. Liu, P. Zhou, Q. Wang, C. Xu, Y. Li, L. Bian, Y. Liu, J. Zhou, F. Wang, Y. Yao, Y. Fang, D. Li, The MTOR signaling pathway regulates macrophage differentiation from mouse myeloid progenitors by inhibiting autophagy. Autophagy 15, 1150–1162 (2019).

25. A. Y. Choo, S. O. Yoon, S. G. Kim, P. P. Roux, J. Blenis, Rapamycin differentially inhibits S6Ks and 4E-BP1 to mediate cell-type-specific repression of mRNA translation. Proc Natl Acad Sci U S A 105, 17414–17419 (2008).

26. D. C. Fingar, J. Blenis, Target of rapamycin (TOR): an integrator of nutrient and growth factor signals and coordinator of cell growth and cell cycle progression. Oncogene 23, 3151–3171 (2004).

27. P. Nicklin, P. Bergman, B. Zhang, E. Triantafellow, H. Wang, B. Nyfeler, H. Yang, M. Hild, C. Kung, C. Wilson, V. E. Myer, J. P. MacKeigan, J. A. Porter, Y. K. Wang, L. C. Cantley, P. M. Finan, L. O. Murphy, Bidirectional transport of amino acids regulates mTOR and autophagy. Cell 136, 521–534 (2009).

28. K. Enomoto, F. Sato, S. Tamagawa, M. Gunduz, N. Onoda, S. Uchino, Y. Muragaki, M. Hotomi, A novel therapeutic approach for anaplastic thyroid cancer through inhibition of LAT1. Sci Rep 9, 14616 (2019).

29. A. L. Haber, M. Biton, N. Rogel, R. H. Herbst, K. Shekhar, C. Smillie, G. Burgin, T. M. Delorey, M. R. Howitt, Y. Katz, I. Tirosh, S. Beyaz, D. Dionne, M. Zhang, R. Raychowdhury, W. S. Garrett, O. Rozenblatt-Rosen, H. N. Shi, O. Yilmaz, R. J. Xavier, A. Regev, A single-cell survey of the small intestinal epithelium. Nature 551, 333–339 (2017).

30. D. Sun, M. Luo, M. Jeong, B. Rodriguez, Z. Xia, R. Hannah, H. Wang, T. Le, K. F. Faull, R. Chen, H. Gu, C. Bock, A. Meissner, B. Gottgens, G. J. Darlington, W. Li, M. A. Goodell, Epigenomic profiling of young and aged HSCs reveals concerted changes during aging that reinforce self-renewal. Cell Stem Cell 14, 673–688 (2014).

31. Y. Zhao, Y. Feng, M. Liu, L. Chen, Q. Meng, X. Tang, S. Wang, L. Liu, L. Li, W. Shen, H. Zhang, Single-cell RNA sequencing analysis reveals alginate oligosaccharides preventing chemotherapy-induced mucositis. Mucosal Immunol 13, 437–448 (2020).

32. D. Li, J. Wang, Ribosome heterogeneity in stem cells and development. J Cell Biol 219, p(2020).

33. R. Y. Ebright, S. Lee, B. S. Wittner, K. L. Niederhoffer, B. T. Nicholson, A. Bardia, S. Truesdell, D. F. Wiley, B. Wesley, S. Li, A. Mai, N. Aceto, N. Vincent-Jordan, A. Szabolcs, B. Chirn, J. Kreuzer, V. Comaills, M. Kalinich, W. Haas, D. T. Ting, M. Toner, S. Vasudevan, D. A. Haber, S. Maheswaran, D. S. Micalizzi, Deregulation of ribosomal protein expression and translation promotes breast cancer metastasis. Science 367, 1468–1473 (2020).

34. K. Kurokawa, Y. Hayakawa, K. Koike, Plasticity of intestinal epithelium: stem cell niches and regulatory signals. International journal of molecular sciences 22, 357 (2020).

35. G. Tomic, E. Morrissey, S. Kozar, S. Ben-Moshe, A. Hoyle, R. Azzarelli, R. Kemp, C. S. R. Chilamakuri, S. Itzkovitz, A. Philpott, D. J. Winton, Phospho-regulation of ATOH1 Is Required for Plasticity of Secretory Progenitors and Tissue Regeneration. Cell Stem Cell 23, 436–443 e437 (2018).

36. F. Ishibashi, H. Shimizu, T. Nakata, S. Fujii, K. Suzuki, A. Kawamoto, S. Anzai, R. Kuno, S. Nagata, G. Ito, T. Murano, T. Mizutani, S. Oshima, K. Tsuchiya, T. Nakamura, M. Watanabe, R. Okamoto, Contribution of ATOH1(+) Cells to the Homeostasis, Repair, and Tumorigenesis of the Colonic Epithelium. Stem Cell Reports 10, 27–42 (2018).

37. A. Ayyaz, S. Kumar, B. Sangiorgi, B. Ghoshal, J. Gosio, S. Ouladan, M. Fink, S. Barutcu, D. Trcka, J. Shen, K. Chan, J. L. Wrana, A. Gregorieff, Single-cell transcriptomes of the regenerating intestine reveal a revival stem cell. Nature 569, 121–125 (2019).

38. K. S. Yan, L. A. Chia, X. Li, A. Ootani, J. Su, J. Y. Lee, N. Su, Y. Luo, S. C. Heilshorn, M. R. Amieva, E. Sangiorgi, M. R. Capecchi, C. J. Kuo, The intestinal stem cell markers Bmi1 and Lgr5 identify two functionally distinct populations. Proc Natl Acad Sci U S A 109, 466–471 (2012).

39. K. Murata, U. Jadhav, S. Madha, J. van Es, J. Dean, A. Cavazza, K. Wucherpfennig, F. Michor, H. Clevers, R. A. Shivdasani, Ascl2-Dependent Cell Dedifferentiation Drives Regeneration of Ablated Intestinal Stem Cells. Cell Stem Cell 26, 377–390 e376 (2020).

40. H. N. Suh, M. J. Kim, Y.-S. Jung, E. M. Lien, S. Jun, J.-I. Park, Quiescence exit of tert+ stem cells by Wnt/β-catenin is indispensable for intestinal regeneration. Cell reports 21, 2571–2584 (2017).

41. A. S. Stewart, C. R. Schaaf, J. A. Luff, J. M. Freund, T. C. Becker, S. R. Tufts, J. B. Robertson, L. M. Gonzalez, HOPX+ injury-resistant intestinal stem cells drive epithelial recovery after severe intestinal ischemia. American Journal of Physiology-Gastrointestinal and Liver Physiology 321, G588–G602 (2021).

42. S. Wirtz, V. Popp, M. Kindermann, K. Gerlach, B. Weigmann, S. Fichtner-Feigl, M. F. Neurath, Chemically induced mouse models of acute and chronic intestinal inflammation. Nat Protoc 12, 1295–1309 (2017).

43. D. A. Guertin, D. M. Sabatini, Defining the role of mTOR in cancer. Cancer Cell 12, 9–22 (2007).

44. D. M. Sabatini, mTOR and cancer: insights into a complex relationship. Nature Reviews Cancer 6, 729–734 (2006).

45. L. Barron, R. C. Sun, B. Aladegbami, C. R. Erwin, B. W. Warner, J. Guo, Intestinal Epithelial-Specific mTORC1 Activation Enhances Intestinal Adaptation After Small Bowel Resection. Cell Mol Gastroenterol Hepatol 3, 231–244 (2017).

46. A. K. Najumudeen, F. Ceteci, S. K. Fey, G. Hamm, R. T. Steven, H. Hall, C. J. Nikula, A. Dexter, T. Murta, A. M. Race, D. Sumpton, N. Vlahov, D. M. Gay, J. R. P. Knight, R. Jackstadt, J. D. G. Leach, R. A. Ridgway, E. R. Johnson, C. Nixon, A. Hedley, K. Gilroy, W. Clark, S. B. Malla, P. D. Dunne, G. Rodriguez-Blanco, S. E. Critchlow, A. Mrowinska, G. Malviya, D. Solovyev, G. Brown, D. Y. Lewis, G. M. Mackay, D. Strathdee, S. Tardito, E. Gottlieb, C. R. G. C. Consortium, Z. Takats, S. T. Barry, R. J. A. Goodwin, J. Bunch, M. Bushell, A. D. Campbell, O. J. Sansom, The amino acid transporter SLC7A5 is required for efficient growth of KRAS-mutant colorectal cancer. Nat Genet 53, 16–26 (2021).

47. S. Yu, K. Tong, Y. Zhao, I. Balasubramanian, G. S. Yap, R. P. Ferraris, E. M. Bonder, M. P. Verzi, N. Gao, Paneth Cell Multipotency Induced by Notch Activation following Injury. Cell Stem Cell 23, 46–59 e45 (2018).

48. Q. Yang, N. A. Bermingham, M. J. Finegold, H. Y. Zoghbi, Requirement of Math1 for secretory cell lineage commitment in the mouse intestine. Science 294, 2155–2158 (2001).

49. R. Ballweg, S. Lee, X. Han, P. K. Maini, H. Byrne, C. I. Hong, T. Zhang, Unraveling the Control of Cell Cycle Periods during Intestinal Stem Cell Differentiation. Biophys J 115, 2250–2258 (2018).

50. T. Sato, J. H. van Es, H. J. Snippert, D. E. Stange, R. G. Vries, M. van den Born, N. Barker, N. F. Shroyer, M. van de Wetering, H. Clevers, Paneth cells constitute the niche for Lgr5 stem cells in intestinal crypts. Nature 469, 415–418 (2011).

51. R. C. Mustata, T. Van Loy, A. Lefort, F. Libert, S. Strollo, G. Vassart, M. I. Garcia, Lgr4 is required for Paneth cell differentiation and maintenance of intestinal stem cells ex vivo. EMBO Rep 12, 558–564 (2011).

52. M. E. Johansson, G. C. Hansson, Mucus and the goblet cell. Digestive diseases 31, 305–309 (2013).

53. H. Jiang, P. H. Patel, A. Kohlmaier, M. O. Grenley, D. G. McEwen, B. A. Edgar, Cytokine/Jak/Stat signaling mediates regeneration and homeostasis in the Drosophila midgut. Cell 137, 1343–1355 (2009).

54. V. Garcia-Hernandez, M. Quiros, A. Nusrat, Intestinal epithelial claudins: expression and regulation in homeostasis and inflammation. Ann N Y Acad Sci 1397, 66–79 (2017).

55. L. Bao, B. Shi, Y. B. Shi, Intestinal homeostasis: a communication between life and death. Cell Biosci 10, 66 (2020).

56. A. Kaser, A. H. Lee, A. Franke, J. N. Glickman, S. Zeissig, H. Tilg, E. E. Nieuwenhuis, D. E. Higgins, S. Schreiber, L. H. Glimcher, R. S. Blumberg, XBP1 links ER stress to intestinal inflammation and confers genetic risk for human inflammatory bowel disease. Cell 134, 743–756 (2008).

57. C. Appenzeller-Herzog, M. N. Hall, Bidirectional crosstalk between endoplasmic reticulum stress and mTOR signaling. Trends Cell Biol 22, 274–282 (2012).

