## supplementary image1-7 for "Amino acid transporter SLC7A5 regulates Paneth cell function to affect the intestinal inflammatory response"

Lingyu Bao *et al.*

**This PDF file includes:**

Figs. S1 to S7
Data S1 to S5(separate files)

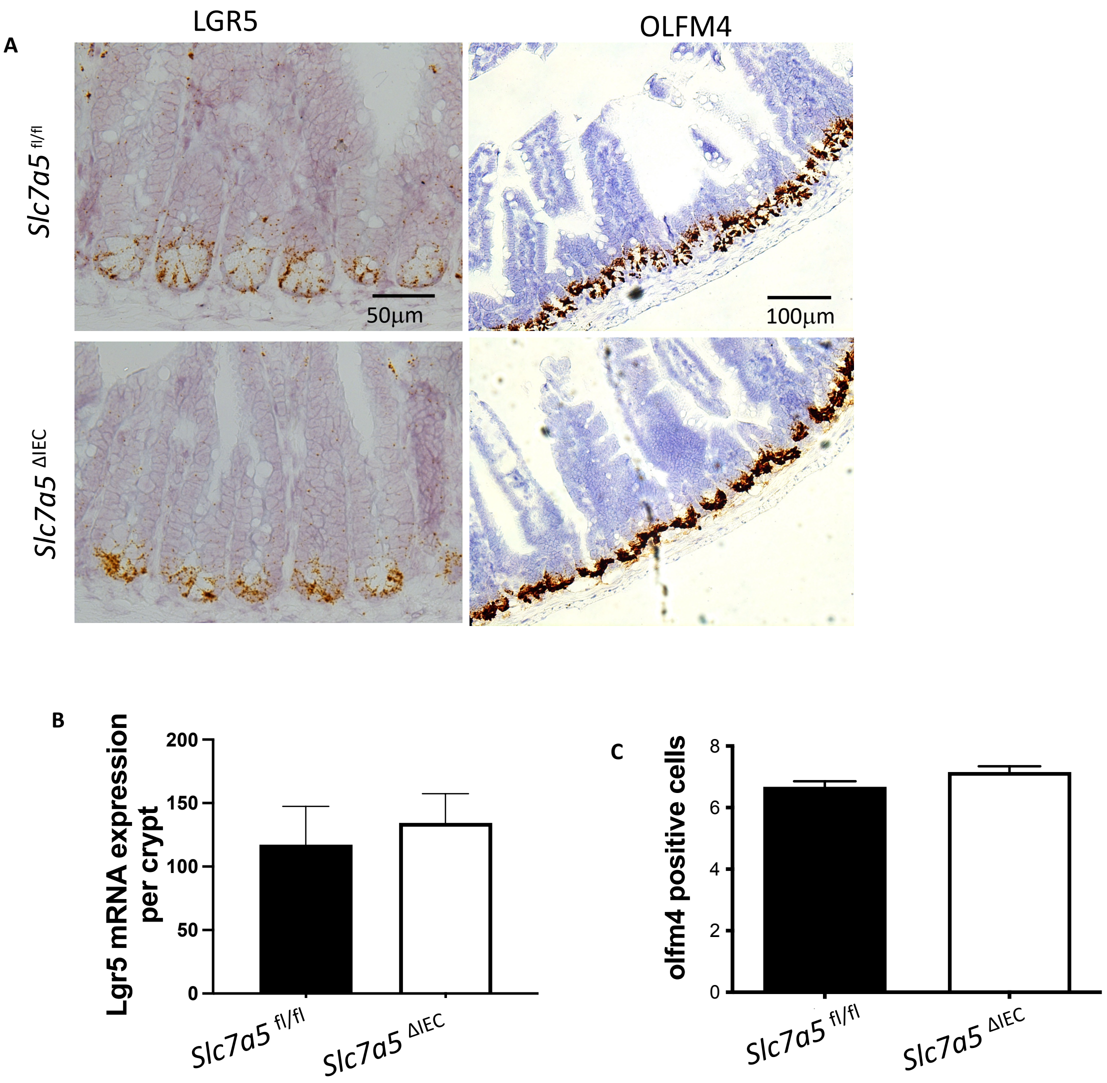


**Fig. S1.** ***Slc7a5*** **$^{\Delta IEC}$ does not affect stem cell marker gene expression in the crypt.**

**A**. Representative pictures of lgr5 and olfm4 single molecule in situ hybridization.

**B-C.** Quantification of the in situ hybridization data showed that LGR5 mRNA level (**B**) and OLFM4+ stem cell numbers (**C**) were similar between *Slc7a5* $^{fl/fl}$ and *Slc7a5* $^{\Delta IEC}$ crypts.

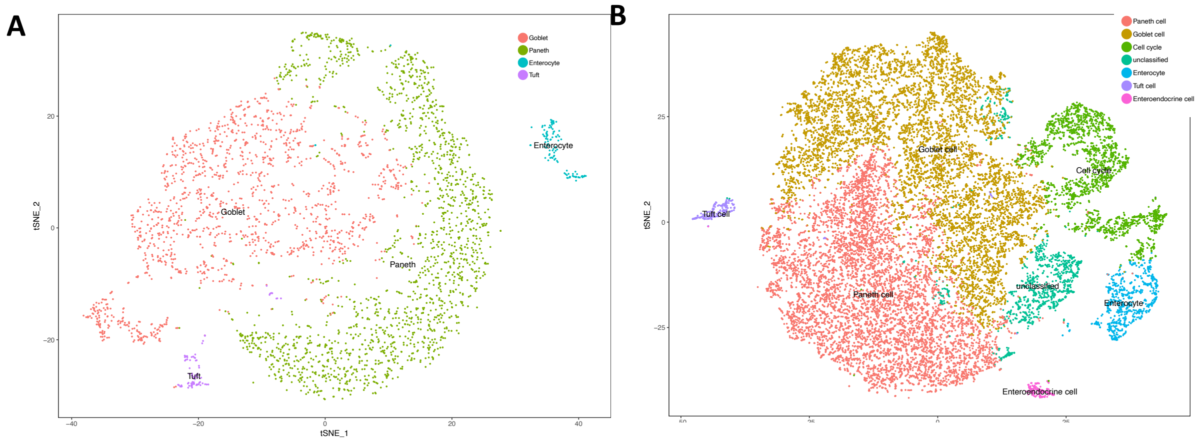


**Fig. S2. ScRNA-seq analysis.** Epithelial cells from intestinal crypts of *slc7a5* $^{fl/fl}$ (A) and *slc7a5* $^{\Delta IEC}$ (B) mice, respectively, were used for scRNA-seq and the cells for each animal type were clustered based on t-SNE plot of the scRNA-seq data with different cell types shown in different colors (note that the cells in cluster labeled as cell cycle were transit amplifying cells or TA cells).

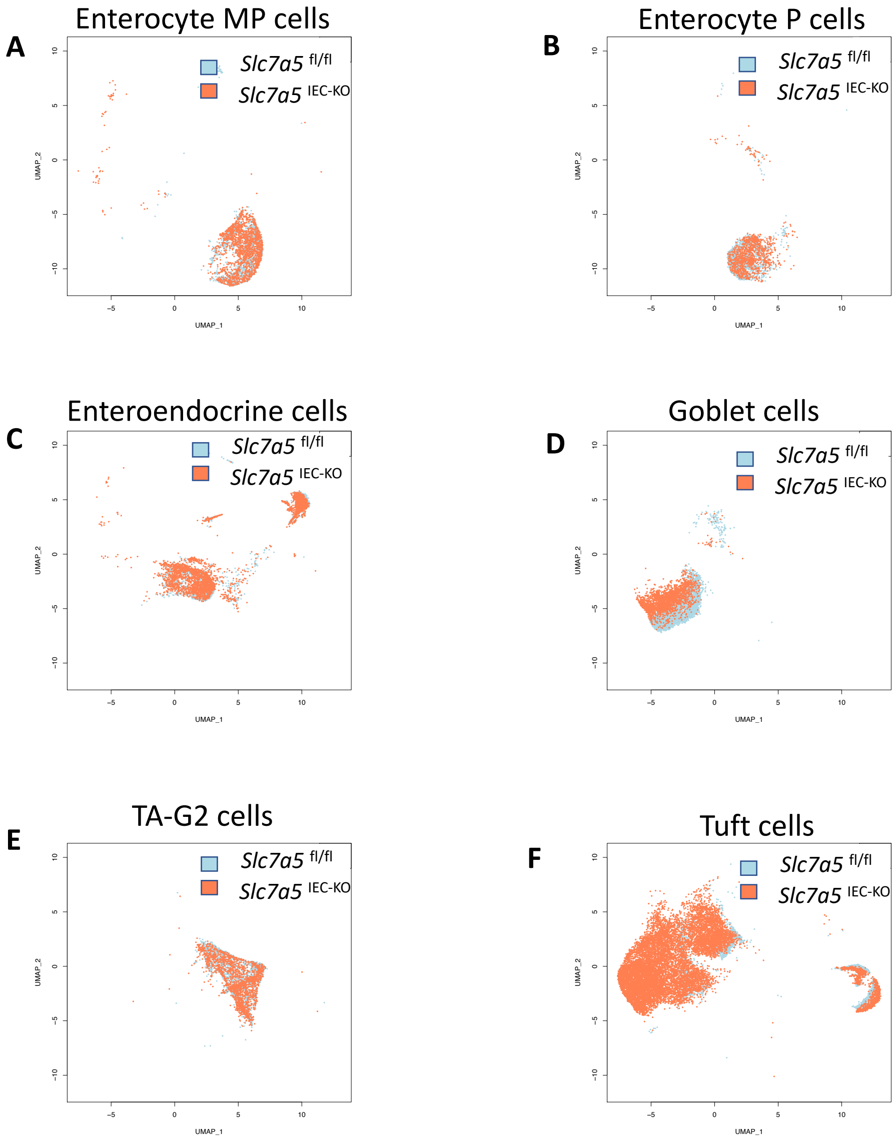


**Fig. S3. Focused views of the regions of the UMAP (Fig. 5C) for the indicated individual epithelial cell types from *Slc7a5* $^{fl/fl}$ mice (blue dots) and *Slc7a5* $^{\Delta IEC}$ (orange dots) mice.** Note that the co-locations of the cells from the *Slc7a5* $^{fl/fl}$ mice and *Slc7a5* $^{\Delta IEC}$ mice for these cell types suggest little or few changes in gene expression between *Slc7a5* $^{fl/fl}$ mice and *Slc7a5* $^{\Delta IEC}$ mice.

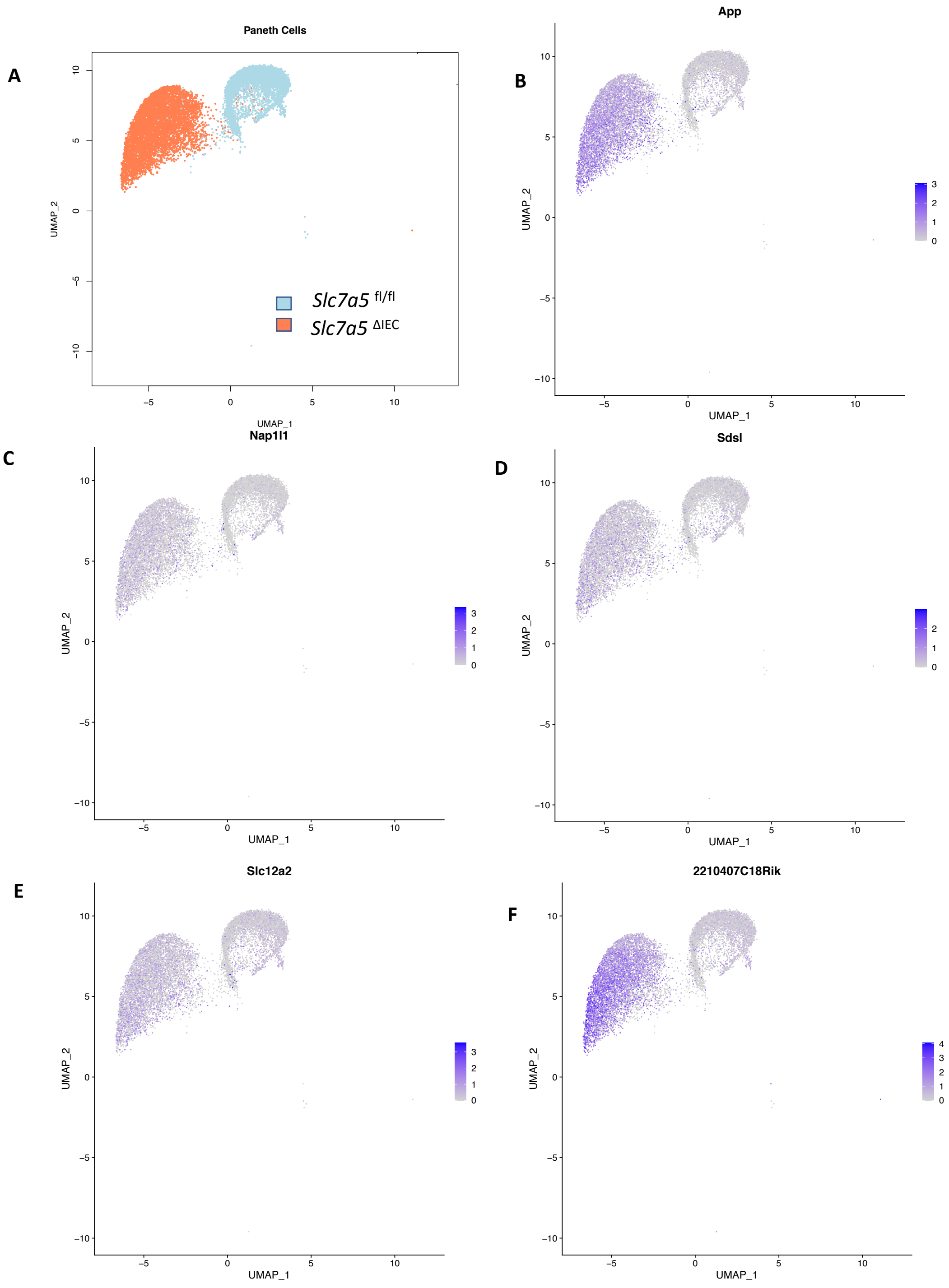


**Fig. S4. UMAP views of increased expression of several stem cell feature genes in the Paneth cells from *Slc7a5* $^{\Delta IEC}$ mice compared to that in *Slc7a5* $^{fl/fl}$ mice.**

**A.** Same image as in figure 5D to show UMAP regions for Paneth cells from *Slc7a5* $^{fl/fl}$ mice (blue dots) and *Slc7a5* $^{IEC\text{-}KO}$ (orange dots) mice.

**B to F:** The expression levels of 5 stem cell feature genes, App, Napl1, Sdsl, slc12a2 and 2210407C18Rik, in Paneth cells as obtained from scRNA-seq were mapped on to the UMAP, showing higher levels in *Slc7a5* $^{\Delta IEC}$ mice than those in *Slc7a5* $^{fl/fl}$ mice.

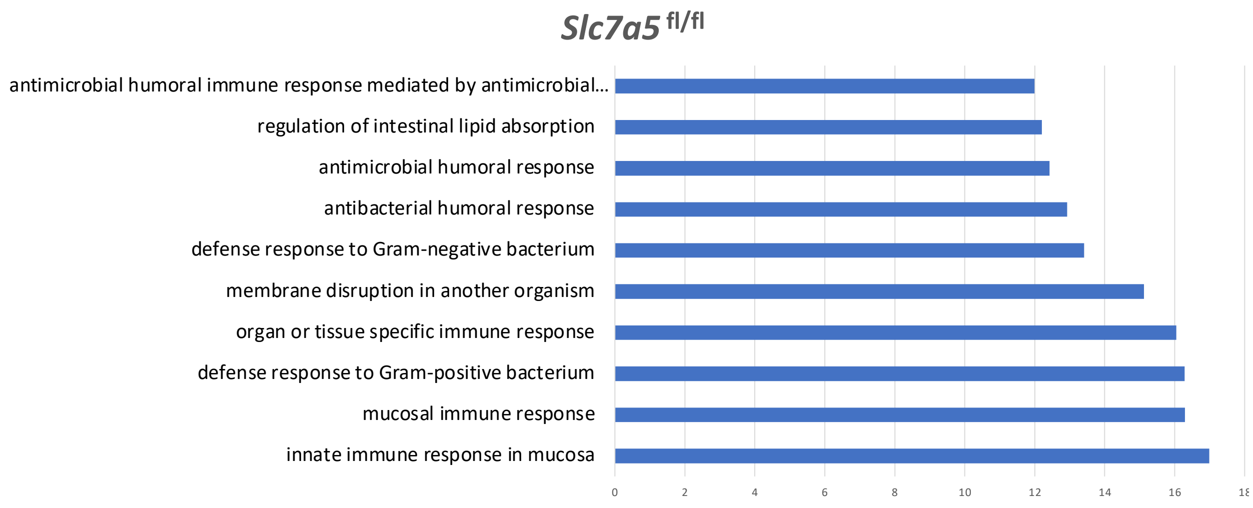


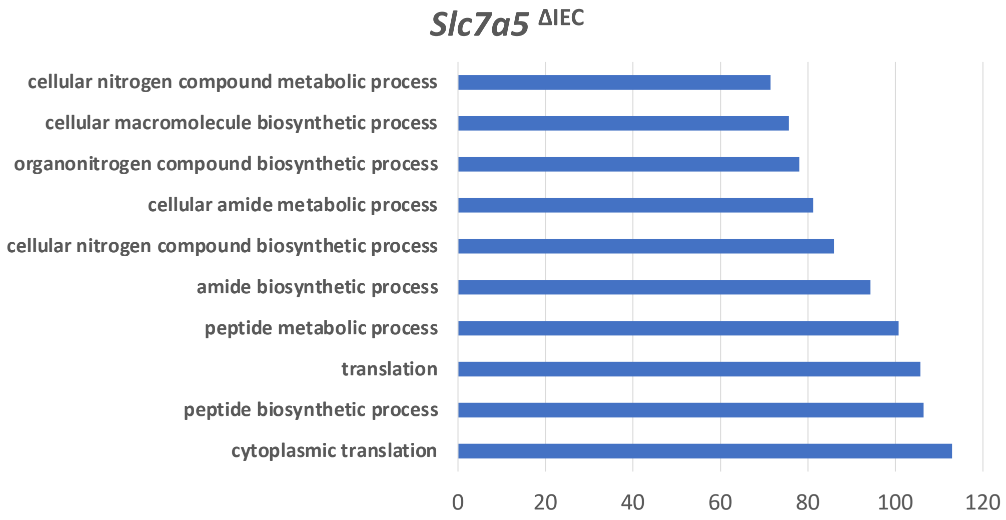


**Fig. S5. GO analysis of DEGs between *Slc7a5* $^{fl/fl}$ mice and *Slc7a5* $^{\Delta IEC}$ mice's Paneth cell.**

**A.** GO terms related to anti-microbial and immune responses were most significantly enriched among the genes expressed at higher levels in the wild type Paneth cells than those in knockout Paneth cells.

**B.** GO terms related to metabolism and biosynthetic processes were most significantly enriched among the genes expressed at higher levels in the knockout Paneth cells than those in wild type Paneth cells.

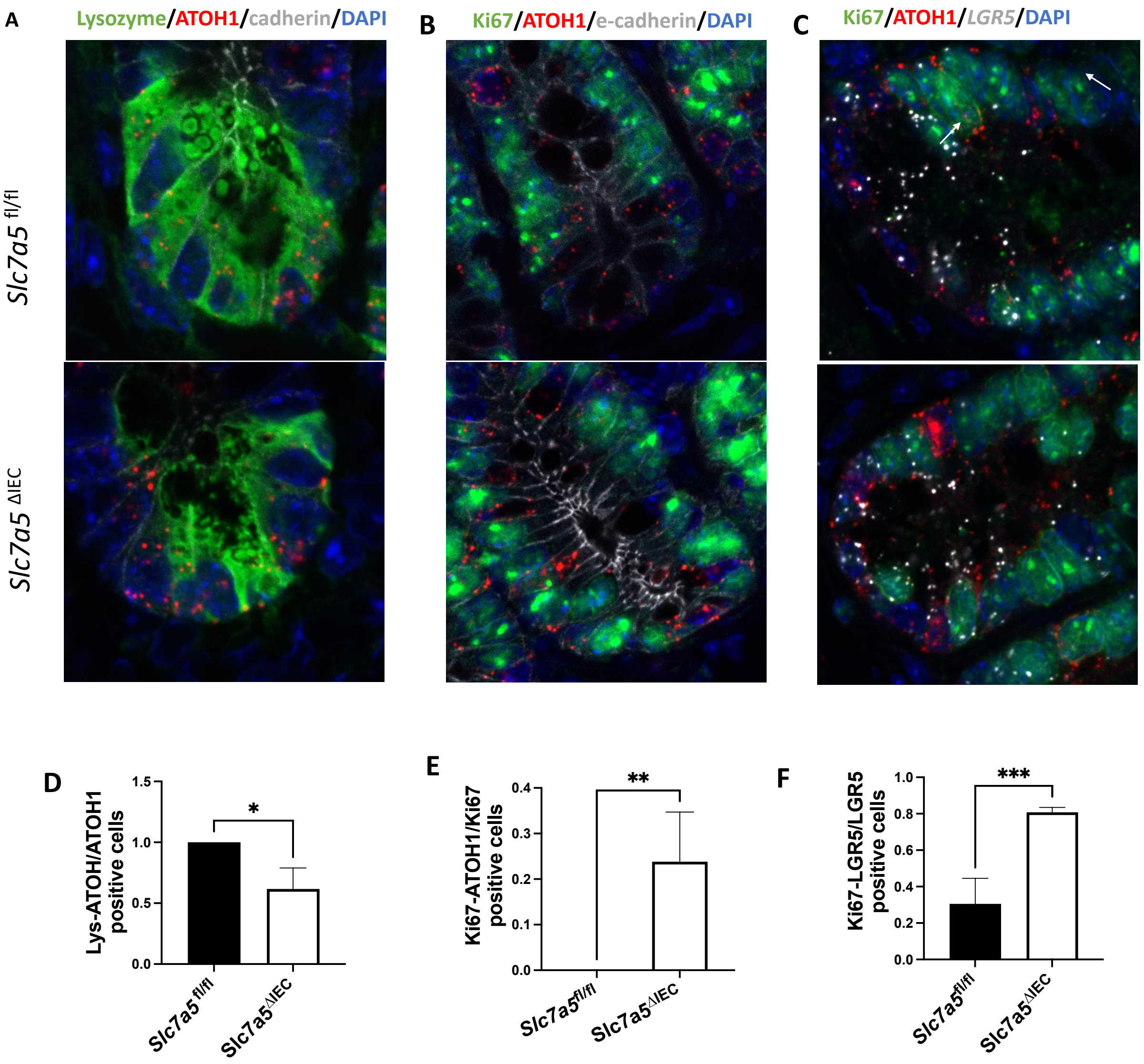


**Fig. S6. Single RNA in situ hybridization and/or immunohistochemical (for lysozyme) analyses of small intestinal crypt base.**

**A**. Representative pictures of lysozyme and atoh1 expression at crypt base in *Slc7a5* $^{fl/fl}$ and *Slc7a5* $^{\Delta IEC}$ mice.

**B.** Representative pictures of Ki67 and atoh1 expression at crypt base in *Slc7a5* $^{fl/fl}$ and *Slc7a5* $^{\Delta IEC}$ mice.

**C.** Representative pictures of Ki67, atoh1, and LGR5 expression at crypt base in *Slc7a5* $^{fl/fl}$ and *Slc7a5* $^{\Delta IEC}$ mice.

**D.** Quantitative analysis of panel A showing that all ATOH1+ cells were lysozyme-positive in *Slc7a5* $^{fl/fl}$ mice, while only about 60% ATOH1+ cells were lysozyme positive in *Slc7a5* $^{\Delta IEC}$ mice.

**E.** Quantitative analysis of panel B showing that Atoh1 were expressed in some proliferating cells in the crypt base of *Slc7a5* $^{\Delta IEC}$ but not *Slc7a5* $^{fl/fl}$ mice.

**F.** Quantitative analysis of panel C showing increased fraction of LGR5+ cells were proliferating in *Slc7a5* $^{\Delta IEC}$ mice crypt base.

| Gene | log2_ratio |
|---|---|
| cxcl1 | 5.26537494 |
| cxcl2 | 4.05088486 |
| Il1rn | 2.92565239 |
| Il1r1 | 2.41463373 |
| cxcl9 | 2.26915173 |
| cxcr2 | 2.00092019 |
| Il11 | 1.47211789 |
| Tnf | 1.34500248 |
| nos2 | 1.28163142 |
| Il17ra | 1.03237594 |
| Il10rb | 1.01003723 |
| Ifngr1 | 0.94563301 |
| Il17rc | 0.90379279 |
| Ilf2 | 0.88726496 |
| Il1rap | 0.87326559 |
| Il4ra | 0.72619279 |

**Fig. S7. IBD (Inflammatory bowel disease)-related genes are upregulated in *Slc7a5* $^{\Delta IEC}$ crypts compared to the wild type crypts.**

The expression levels for these genes based on the total normalized gene counts from scRNA-seq in the knockout crypts were divided by the corresponding values in the wild type crypts and the ratio for each gene was presented in log2 value.

**Data S1. DEGs upregulated in Paneth cells from *Slc7a5* $^{fl/fl}$ mice compared *Slc7a5* $^{\Delta IEC}$ mice**

**Data S2. DEGs upregulated in Paneth cells from *Slc7a5* $^{\Delta IEC}$ mice compared *Slc7a5* $^{fl/fl}$ mice**

**Data S3. GO terms enriched among DEGs expressed at higher levels in the Paneth cells from *Slc7a5* $^{fl/fl}$ mice**

**Data S4. GO terms enriched among DEGs expressed at higher levels in the Paneth cells from *Slc7a5* $^{\Delta IEC}$ mice**

**Data S5. The expression of inflammatory gene (normalized counts from scRNA-seq) in *Slc7a5* $^{fl/fl}$ and slc7a5 $^{\Delta IEC}$ mice**
